# Neural systems for memory-based value judgment and decision-making

**DOI:** 10.1101/712661

**Authors:** Avinash R. Vaidya, David Badre

**Author notes:** Address correspondence to: Avinash R. Vaidya, Brown University, Providence, RI 02912-1978.

## Abstract

Real life choices often require that we draw inferences about the value of options based on structured, schematic knowledge about their utility for our current goals. Other times, value information may be retrieved directly from a specific prior experience with an option. In a functional magnetic resonance imaging (fMRI) experiment, we investigated the neural systems involved in retrieving and assessing information from different memory sources to support value-based choice. Participants completed a task in which items could be conferred positive or negative value based on schematic associations (i.e. schema value), or learned directly from experience via deterministic feedback (i.e. experienced value). We found that ventromedial prefrontal cortex (vmPFC) activity correlated with the influence of both experience- and schema-based values on participants’ decisions. Connectivity between vmPFC and middle temporal cortex also tracked the inferred value of items based on schematic associations on the first presentation of ingredients, prior to any feedback. In contrast, the striatum responded to participants’ willingness to bet on ingredients as a function of the unsigned strength of their memory for those options’ values. These results argue that striatum and vmPFC play distinct roles in memory-based value judgment and decision-making. Specifically, the vmPFC assesses the value of options based on information inferred from schematic knowledge and retrieved from prior direct experience, while the striatum controls a decision to act on options based on memory strength.

## Introduction

How can we predict the value of options without prior, explicit experience? For example, when building a table, a hammer has greater value than a ball of yarn. No previous experience using yarn in place of a hammer is needed to make this kind of value judgment. However, this evaluation does depend on knowledge about yarn, hammers and table construction. Structured knowledge about the world that can support this kind of inference is often referred to as a schema (Bartlett, 1932; Ghosh & Gilboa, 2014; Piaget, 1952). Judging the value of options for accomplishing complex goals, like table building, requires inference based on these schematic representations.

Value does not always need to be inferred. In the same example, seeing a certain brand of varnish on display might remind you that your friend had recommended that varnish for your table last week. In this case, value relies on the retrieval of an explicit positive association with the item from a specific time and place. Here, value is a memorized feature of one or a few previous episodes, which may be brought to mind through mechanisms of episodic retrieval.

Relatively little is known about the neural systems involved in retrieving information from different memory sources to support value judgment. Lesion studies have shown that ventromedial prefrontal cortex (vmPFC) and orbitofrontal cortex (OFC) are necessary for decision-making when option values must be inferred from knowledge of task structure, but not when values can be retrieved from directly learned associations (Bradfield, Dezfouli, van Holstein, Chieng, & Balleine, 2015; Izquierdo, Suda, & Murray, 2004; Jones et al., 2012; Reber et al., 2017). It is unclear if these deficits arise because these regions represent task structure (Schuck, Cai, Wilson, & Niv, 2016; Wilson, Takahashi, Schoenbaum, & Niv, 2014), guide retrieval based on these schemas (Eichenbaum, 2017), or specially utilize structured knowledge during value judgment.

A parallel line of research has focused on the striatum at the intersection of goal-directed retrieval and value-based decision-making. Recent work has argued that the striatum is involved in assessing the value of information stored in memory and using this information to bias cortical representations of choices with higher value (Shadlen & Shohamy, 2016). A mutually compatible alternative is that striatum integrates memory signals into its basic decision making functions, including those related to memory itself (Han, Huettel, Raposo, Adcock, & Dobbins, 2010; Scimeca & Badre, 2012), though these may also be separable processes (Elward, Vilberg, & Rugg, 2015).

The function of these regions in value judgment likely further depends on coordination with several other regions for the retrieval of relevant information. Notably, overlapping regions within the vmPFC and the striatum are connected with regions canonically associated with episodic and schematic memory, and adaptive retrieval, including the hippocampus (HPC), middle temporal cortex (MTC) and anterior ventrolateral prefrontal cortex (aVLPFC) (Haber & Knutson, 2010; Price, 2007). However, it is not known how these regions interact to link retrieval to decision-making.

We tested three hypotheses from this prior literature on memory and value-based decision making using a functional magnetic resonance imaging (fMRI) experiment: (1) vmPFC is preferentially engaged when value judgments are based on structured knowledge about the world (i.e. schema memory). (2) Striatum preferentially tracks experienced values, in keeping with its role in both episodic memory and valuation. (3) vmPFC and striatum interact with different memory stores in accord with their respective roles in constructing values through schema-based inference (vmPFC) versus direct, explicit experience (striatum).

## Methods

### Behavioral Task and Experimental design

We investigated the neural systems involved in making value judgments based on different memory sources using a novel experimental task. This task, and all other tasks not administered through Amazon Mechanical Turk (AMT), were programmed in Psychtoolbox-3 (Brainard, 1997; Kleiner, Brainard, & Pelli, 2007). In this experiment, participants took the role of a restaurant chef choosing whether or not to serve ingredients to a customer (Figure 1). Each trial required participants to make a bet based on their belief that the customer would like or dislike the displayed ingredient. Participants were instructed that if they served an ingredient that the customer liked, they would receive 50 points, however if they served an ingredient that the customer did not like, they would lose 25 points. If participants chose not to serve an ingredient, they would receive 0 points. This asymmetry in the points rewarded versus lost for betting trials was chosen to offset the overall loss aversion of participants. In this task, loss aversion is expressed as a tendency to avoid endorsing ingredients on the first presentation, as revealed in behavioral piloting.

**Figure 1.**
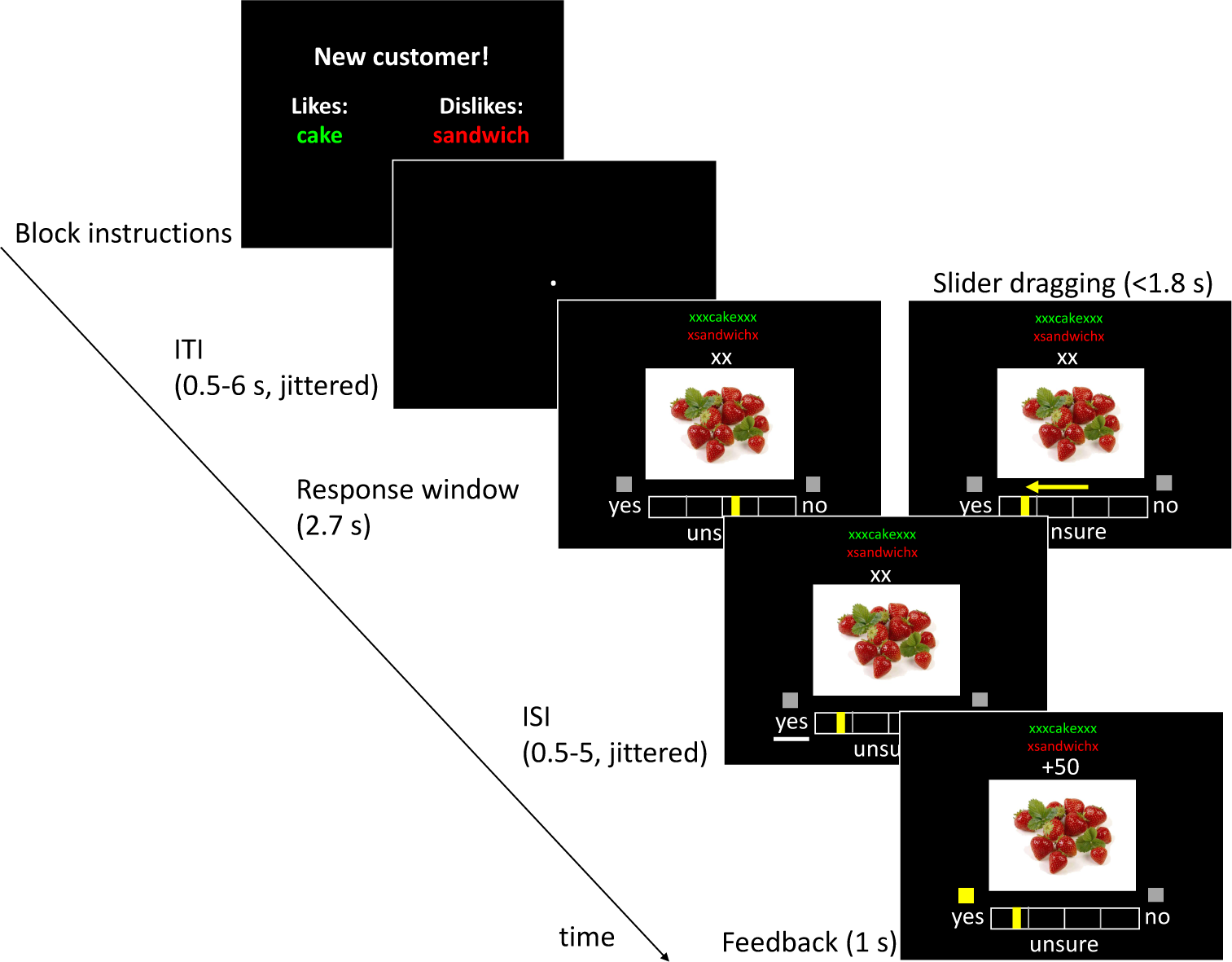
Task schematic showing instructions at the start of a block and organization of a single example trial. At the start of the block, participants were informed of the recipes that customer liked and disliked. These recipe names were presented in green and red above the ingredient on all trials. In each trial, participants indicated their decision to feed the displayed ingredient to the current customer, and their confidence in this decision, using a slider. Participants then saw whether or not the customer liked the ingredient, signaled by a box above either response cue, and the points earned or lost in each trial. ITI (inter-trial interval), ISI (inter-stimulus interval).

After the participant made their choice, the customer’s preference for a given ingredient was always signaled by a cue above the correct response during feedback, regardless of whether the participant chose to feed the customer or not, and all customer preferences were deterministic. Thus, participants received complete information on each trial regardless of their response. The feedback provided a source for learning the value of an individual ingredient. If the participant did not respond in time, no information about the customer’s preferences was given and participants were shown the word ‘slow’ in place of feedback about points earned.

In addition to the information about ingredient preferences provided through feedback, participants were told at the start of a block that the customer liked ingredients pertaining to one recipe (positive recipe) and disliked another (negative recipe). In this way, the participant could also infer the value of an ingredient on the basis of schematic knowledge about what ingredients go with what recipes, without having received feedback. For example, if the customer liked cake and disliked soup, flour might be positively valued and onions might be negative by virtue of their association with each recipe. Participants were instructed that these recipes were general and could encompass many potential variants (e.g. the recipe ‘tacos’ might include both beef and fish tacos). The names of these recipes were displayed at the top of the screen in each trial, with the positive recipe shown in green and the negative recipe in red text.

At the start of each block, participants were told that they were about to feed a new customer, and were given the names of the recipes that the customer liked and disliked. The two recipes in each block were randomly selected without replacement from a set of six recipes. The screen was then left blank for 12 s until the start of the first trial.

Participants were also instructed before the start of the task that each customer liked and disliked other ingredients that were unrelated to these recipes (i.e. ‘independent ingredients’). The value related to these ingredients could not be anticipated in advance based on their schematic associations, but rather would have to be acquired through experience.

Each ingredient was repeated three times within each block. For these repeated ingredients, participants could retrieve a memory of previous feedback to make a decision. All repeated presentations were separated by at least one other ingredient so that participants were never making decisions about the same ingredient for multiple trials successively.

Thus, the experiment had three factors by design: ingredient clustering (recipe or independent), value (positive or negative) and presentation number (1, 2, 3). For the purposes of behavioral and fMRI analysis, any missed responses were not considered in the count of ingredient presentations, as no value information was communicated to the participant on these trials (e.g. if a participant missed the response deadline on the first presentation, the second presentation would be considered the first, and the third presentation would be considered the second in all analyses).

In each trial, participants saw an ingredient and were asked to respond by moving a slider to either side of a bar to indicate their decision to serve an ingredient and their confidence in that decision with a single response, similar to Scimeca, Katzman, and Badre (2016). Moving the slider past the midway point to either side was considered a binary selection of that side’s response, while the distance of the slider from the midway point provided a continuous index of confidence. The slider would start in a random position on each trial and participants could then drag it toward the left or the right by holding down a button on a fiber optic response pad (Current Designs, Philadelphia, PA) using their right index or middle fingers. The slider would stop moving when it reached an end of the bar, or when participants let the button go. Participants could not change the slider position after letting go of the button. The side of bar where participants placed the slider signaled their choice, and the distance of the from the middle of the bar indicated their confidence.

Before the fMRI experiment, participants completed a practice task in which they learned to use the slider while verifying various general knowledge statements (e.g., ‘Black bears are omnivores’), and indicate their confidence in their response. Participants received further practice outside the scanner by completing a short practice block of the restaurant task where they fed a customer with recipes and ingredients that did not appear in the fMRI experiment. Before completing this practice block, all participants had to provide a detailed verbal description of the task instructions, including the payoff structure, to the experimenter. The experimenter verbally filled in any gaps in the participants’ explanation of the task before allowing participants to proceed into the practice phase.

Trials were preceded by a jittered inter-trial interval (ITI) drawn from a lognormal distribution (0.5-6 s). In each trial, participants saw a food ingredient centrally displayed for up to 2.7 s, during which time they could choose whether or not to feed the ingredient to the customer (‘yes’/’no’). Stimulus presentations duration did not depend on response time. The names of the recipes that each customer liked and disliked appeared at the top of the screen on each trial, color coded to indicate the customer’s preference.

After a response, there was a jittered inter-stimulus interval (ISI) drawn from a uniform distribution (0.5-5.5 s). The remaining time from the response window and slider movement period was also added to this interval. Participants then saw feedback about their decision for 1 s. A square above the correct answer would light up, and the amount of points added or subtracted from participants’ total would be displayed in this period.

Trial-order, ITI and ISI lengths were drawn from sequences optimized for efficient estimation of the hemodynamic response across conditions of interest. These sequences were optimized by generating 5000 random sequences, and using SPM12 to create event-based design matrices for stimuli and feedback events in each sequence. The average variance for each regressor explained by all other regressors in the matrix was calculated for each sequence and multiplied by the total duration of the sequence to approximate design efficiency. The three sequences with the lowest values in this measure (i.e. least covariance in regressors with the shortest experiment time) were used for the fMRI experiment and were randomly ordered for each subject. Participants completed three blocks of this task in the scanner, each a separate fMRI run. Each block consisted of 84 trials total, with seven trials for each value-clustering-presentation condition in each block (21 trials per condition total).

### Participants

Twenty-six participants were recruited from the Providence, RI area to participate in the fMRI experiment. Two participants were excluded due to movement greater than our voxel size on at least one dimension during the functional scan, and one was excluded for falling asleep during the experiment. Of the remaining 23 participants, four were male, their mean age was 22.5 years (SD = 4.0), and their mean years of education was 15.5 (SD = 2.3). All participants were right-handed, had normal or corrected-to-normal vision, and were screened for the presence of psychiatric or neurological conditions, psychoactive medication use, and contraindications for MRI. Two participants rejected all ingredients on the first presentation of the positive independent condition, prohibiting analysis of their behavior or ROI-level responses in a full 12-condition ANOVA after filtering for correct responses. One participant did not commit any errors on the second or third presentation of the positive recipe ingredients, prohibiting analysis of their error trials in this condition.

Two hundred participants from the United States were recruited through Amazon Mechanical Turk (AMT) to provide normative naming responses for images of food ingredients. Four of these subjects were rejected for exceptionally low accuracy in this task (< 50%). The mean age of the remaining participants was 35.4 years (SD = 10.5), and 107 of them were male.

A second sample of 221 participants were recruited from the United States through AMT to provide normative data for ingredient-recipe associations. Twenty-nine of these participants were rejected for failing to follow task instructions (most often by simply writing the names of the ingredients rather than associated recipes). The remaining participants had a mean age of 36.0 years (SD = 11.1), and 71 of them were male.

All participants across experimental and norming studies gave informed consent as approved by the Human Research Protections Office at Brown University, and were compensated for their participation. Participants in the fMRI experiment also received a monetary bonus proportional to the points they earned in the main task.

### Stimuli

Ingredient stimuli were obtained from the bank of standardized stimuli (BOSS; N = 65) (Brodeur, Dionne-Dostie, Montreuil, & Lepage, 2010; Brodeur, Guerard, & Bouras, 2014), food-pics database (N = 50) (Blechert, Meule, Busch, & Ohla, 2014), with a small set of supplemental food images collected by A.V. from Google Images (N = 4). These images were selected to include basic food ingredients — i.e. items that might be used as parts of a recipe, rather than completed meals. For example, an image of cheese would be included rather than an image of a pizza. All images were scaled to the same size and padded with white pixels or cropped to fit the same area. Ingredients from the BOSS database were excluded if more than 20% of the normative sample either responded that they did not know the name of the ingredient or did not recognize the ingredient. Ingredients were also rejected if the modal name for the ingredient given by respondents was incorrect (indicating low recognition accuracy), and if the H-value (a measure of naming disagreement across participants, similar to entropy) was above 3.00.

Because similar naming data were not available for ingredients from food-pics or Google Images, normative naming data were obtained through an AMT experiment. Participants were shown ingredient images and asked to write the name of the ingredient in a text box below, or write “I don’t know.” Responses that included common variants of ingredient names were considered correct, while overly general names were marked as incorrect (e.g. ‘nuts’ for an image of pecans). Ingredients where more than 20% of participants did not know the name, or gave incorrect answers, were rejected. Naming agreement was also examined by calculating an H-value for each ingredient; however, this value did not exceed 3.00 for any ingredient in this set.

### Normative recipe associations

Associations between ingredients and recipes were obtained through a norming experiment conducted on AMT akin to single word generation paradigms used to derive word associations (Postman & Keppel, 1970). Participants were shown ingredient images one at a time and were asked to write the name of the first recipe that came to mind using the ingredient shown on screen. Unique recipe names were then extracted for each ingredient and two researchers (A.V. and H.J.) manually processed these data. Alternative names, semantically similar responses and misspellings were all collapsed into single recipes, and ingredient specific information was removed to facilitate detection of overlap in ingredient-recipe associations (e.g. responses like ‘pumpkin soup’ and ‘mushroom soup’ were both reclassified as ‘soup’). The end result of this process was a matrix with endorsement counts for 102 ingredients in 1876 recipes. Six ingredients were then dropped from this set, as they were visually too similar to other ingredients in the experiment, leaving 96 ingredients total.

Of these recipes, many (915) were only named by a single AMT participant for a single ingredient. As our task design required that each recipe have a large number of associate ingredients (at least seven recipe ingredients, and some additional ingredients to be used as independent items), we cycled through different parameters for filtering out recipes. With each of these filtered lists, ingredients were assigned to hierarchically organized clusters using the Matlab (Mathworks, Natick, MA) function ‘linkage,’ which sorted ingredients based on the mean cosine similarity of their naming frequencies. Through this process we selected six recipes where the clustering solution provided ingredient cluster sizes that were large enough for our task design (pasta (N=10), soup (N=13), cake (N=15), pie (N=21), salad (N=17) and sandwich (N=20)).

### Value rating task

At the start of the experimental session, participants completed a short rating task where they were asked to provide ratings for how much they would hypothetically want to have each ingredient at the end of the session. Participants provided responses on a seven point Likert scale from −3 to 3, with labels on −3 reading ‘Not at all,’ 0 reading ‘Indifferent’ and 3 reading ‘Very much.’ These subjective value ratings were included as a nuisance variable in each of the parametric GLM analyses, and used to test if participants’ personal preferences influenced their decisions in the task in model-based analyses. This task also served to visually familiarize participants with all ingredient images.

### Computational modelling of behavior

Computational models of behavior were used to characterize the dynamics underlying participants’ behavior in this task. We developed a simple base model that made a set of assumptions about behavior in this task that were carried over to all alternative models. Each alternative model made a modification of this base model, or the best model from the last step. Alternative model modifications were assessed and compared using a cross-validation procedure, described in detail below. For simplicity, we present the final best fitting model that resulted from this process. We also briefly describe alternative model modifications that did not survive the selection process in the following section.

Within the best fitting model, the value (*V)* of the ingredient (*i)* on each trial was calculated as:

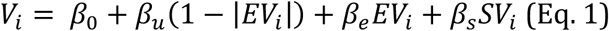

Where *β*_*e*_ controls the influence of experienced values (*EV*), and *β*_*s*_ controls the influence of schema values (*SV*) for the current trial ingredient (these two terms were included in the base model). The parameter *β*_*u*_ is a free temperature parameter that controls the model’s tendency to endorse or reject an ingredient based on its unfamiliarity (the opposite of the absolute value of the experienced value term — described below; unfamiliarity model). An intercept term, *β*_0_ captures overall, item-independent biases toward betting or passing (fixed bias). The model’s willingness to bet on each trial (i.e. the model-P(yes)) was then calculated by placing *V*_*i*_ in a simple logistic function.

On each trial where participants responded, the experienced value for each ingredient was updated according to a simple Rescorla-Wagner learning rule (Rescorla & Wagner, 1972):

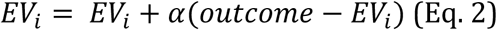

Where the value for *outcome* could be 1 or −1, depending on whether the feedback was positive or negative. While this rule is more commonly used in the context of reinforcement learning, it was adopted here as a simple means of operationalizing a graded memory that is signed and becomes stronger with repetitions of consistent value associations. The parameter *α* was a fixed learning rate that was fit for each subject in an alternate version of the base model where *β*_*e*_ was held at different constant values (1, 2, 5 and 8). The model fit with a constant value of 5 for *β*_*e*_had the lowest average negative log-likelihood of constants tested, and so *α* values from this fit were used in the base model and all further variations. This procedure to separately fit *α* was followed because *β*_*e*_ and *α* could not be stably estimated within the same model. Estimating these parameters separately allowed us to roughly capture individual differences in the encoding of *EV,* while separately estimating the contribution of *EV* to choice behavior through *β*_*e*_. Allowing *α* to vary was also found to improve the fitness of the base model over a variant where *α* was fixed to 1.0.

To reflect forgetting, the values for all ingredients also decayed towards zero with each trial according to:

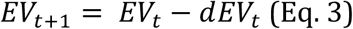

Where *d* is a parameter that controls the rate of this decay (also included in the base model).

Schema values were calculated through a weighted difference of the associative strength of ingredients with positive and negative recipes. The associative strength of each ingredient with these recipes was determined through a procedure based on latent semantic analysis (LSA) (Dumais, 2004; Landauer, Foltz, & Laham, 1998). The endorsement frequencies for ingredients across all recipes in the normative AMT data were weighted by their entropy across recipes and a singular value decomposition (SVD) was carried out on this matrix. This matrix was then reconstituted with a smaller number of dimensions set by a free parameter *c* (dimensions model). This parameter essentially controlled the dimensionality of recipe space that the model would consider, so that a lower dimensional model would separate ingredients on only simple features (i.e. savory or sweet), while a higher dimensional model would include increasingly granular information that separates ingredients along information about specific recipes (Figure 2). Notably, allowing this parameter to vary did not dramatically influence the overall choice behavior of the model in simulations, but did change the shape of schema values so that increased dimensionality increased the discretization within this term. The association of an ingredient with each recipe was calculated as the mean cosine similarity between that ingredient and all ingredients in each recipe cluster within this reconstituted matrix. The schema value for each ingredient was then calculated as the difference in the associative strength for the positive and negative recipe.

**Figure 2.**
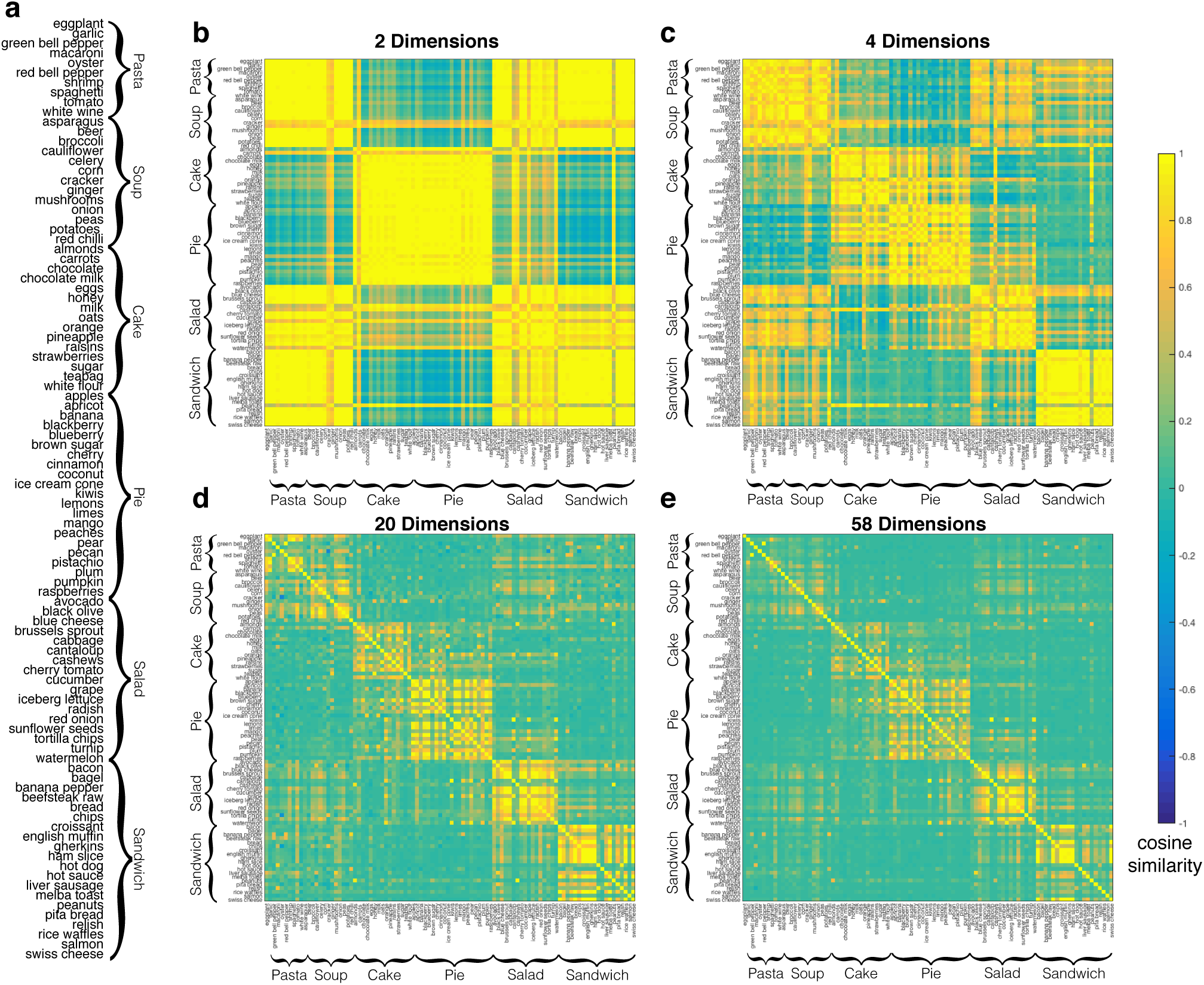
Example ingredient by ingredient cosine similarity matrices from reconstructions of ingredient-recipe associations with increasing dimensions. **a.** Ingredient names and clustering by recipe. **b.** Cosine similarity matrix with two dimensions **c.** with four dimensions, **d.** with 20 dimensions, and **e.** with 58 dimensions.

In subsequent presentations, the model discounted schema values with an exponential function based on memory strength for experienced values (i.e. the absolute value of *EV*):

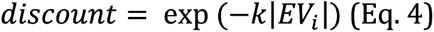

Where *k* was a free parameter that determined the steepness of this discounting. This discount factor was then included in the calculation of ingredient values:

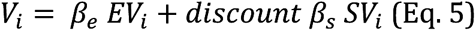

So that the contribution of schema values scaled with this exponential discounting factor.

### Alternate model parameters

Briefly, the following alternate modifications to the base model were considered, but did not improve model fitness:

#### (1) Asymmetric schemas model

This model included separate temperature parameters for positive and negative schema values, so that the value of an ingredient was:

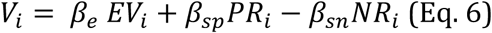

Where *β*_*sp*_ and *β*_*sn*_ are temperature parameters that control the influence of ingredient associations with positive and negative recipes, respectively.

#### (2) Schema similarity model

The recipes in a block could vary in their similarity, making some ingredients plausibly associated with both positive or negative recipes (e.g. pie and cake). To accommodate this possibility, we tested a model where the associative strength of ingredients with the positive and negative schemas to the schema was weighted by an estimate of participants’ uncertainty about the recipe associations of each ingredient, calculated as the probability that an ingredient was associated with both recipes (schema similarity model; P(*SR*)):

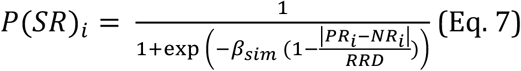

Where *RRD* corresponds to the range of recipe association differences (i.e. the difference between the maximum and minimum recipe associative strength for all ingredients and recipes). This probability was then used in calculating weights for the sum and difference of the positive and negative recipe associations, where the weight for the sum of these associations was calculated as

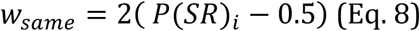

And the weight for the difference of these ingredient associations was calculated as

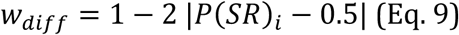

*SV* was then calculated as the weighted combination of the difference and sum of each ingredient’s associative strength with the positive and negative recipes:

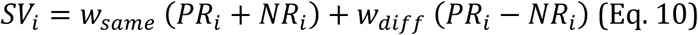

The parameter *β*_*sim*_ controlled sensitivity to the similarity of ingredients’ recipe associations, and the relative contribution of the difference or sum of positive and recipe associations to *SV*. This parameter also controlled the valence of the additive strength of these associations: positive or negative values for *β*_*sim*_ would give the same valence to the additive strength of recipe associations, respectively. Alternatively, if the *β*_*sim*_ parameter was zero, participants would rely solely on the difference of the ingredient’s associative strength with the positive and negative recipes.

#### (3) Dual process recollection model

While the above models assume that experienced values are stored in episodic memory via a graded process, other models developed in memory judgment tasks suggest that episodic memory has a graded component (familiarity) and a discrete component (recollection) (Yonelinas, 1994). To test whether experienced values in this task may also rely on an additional non-graded recollection process, ingredient values were calculated as:

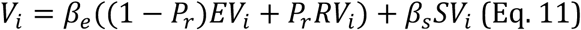

Where *RV* is the recollected value of an ingredient, i.e. the outcome from previous trials that does not decay or change over time (−1 or 1), and *P*_*r*_ is a fixed probability of recollection that does not change over trials. Thus, in this model, values learned through feedback contributed to the decision as the sum of recalled and graded value information, weighted by the probability of recall.

#### (4) Schema spreading model

<u>Rather than treat experienced and schema-based values separately, it is possible that participants may have used schematic knowledge to try and facilitate feedback-based learning. To test this hypothesis, we created a model where values learned through feedback were broadcast to other ingredients, weighted by their similarity in recipe space. Upon feedback, the experienced value for ingredient</u> *<u>i</u>* <u>was used to update a schema spread value term (SSV) for all other ingredients (</u>*<u>j</u>*<u>) according to:</u>

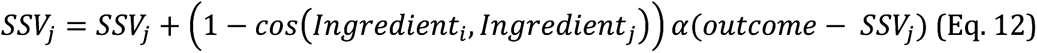

<u>Where</u> *α* is the same learning rate used for updating experienced values and *cos* refers to the cosine distance between ingredients in the normative ingredient-recipe association matrix produced through LSA.

This SSV term was included in calculating the value of each ingredient *i* in each trial, weighted by an additional free temperature parameter.

(5) Participant subjective value

We also tested an alternate model that included participants’ own subjective value ratings for ingredients in the computation of ingredient values weighted by an additional free temperature parameter. This model tested whether participants’ choices would be better accounted for by taking into account their personal preferences.

### Model fitting and comparison

Participants’ responses consisted of two main features: a binary response indicating whether or not to feed an ingredient to a customer, and a continuous rating of their confidence in this binary response. This confidence response could also be transformed into a continuous measure of participants’ belief that a customer would like an ingredient (i.e. P(yes)) by measuring the position of the slider between the ‘yes’ and ‘no’ options rather than the distance of the slider from the middle, similar to Scimeca et al. (2016). Models were fit to both participants’ continuous and binary choice responses. The overall model likelihood was calculated as the product of the model likelihood for participants’ binary responses, and the model likelihood of the linear fit between the logit transformed model-P(yes) and participant-P(yes). The negative log of this likelihood function served as the objective function to be minimized in the fitting procedure carried out by the Matlab function ‘fmincon.’

Computational models were fit to participants’ behavior in a step-wise process. At each step, a simpler preferred model was modified by the addition of new free parameters. Models were compared with a cross-validation procedure. For each participant, we generated 100 independent training and test datasets where a set of 16 ingredients (19% of the total ingredients) were randomly held out of the data used for fitting model parameters and treated as deadline failures (as to maintain the same number of trials in the experiment). We chose to randomly subsample ingredients rather than blocks or trials because of the structure of the task: participants encounter a new customer, and thus a new set of recipe schemas in each block, while trials are not independent from each other, as each ingredient is repeated three times and the value must be retained from one trial to the next.

To test model fitness, we estimated the mean difference in the negative log-likelihood of each model relative to the base model for the held out 16 ingredients, averaged across all 100 test datasets and summed across participants. At each step in the model comparison process, the model that resulted in the largest reduction in the negative log-likelihood for the test data set was considered the preferred model. Additional parameters reduced the negative log-likelihood were added to the preferred model and compared in the next step, while any parameters that did not improve model fitness were discarded in future steps. The search for the best fitting model was stopped once additional parameters no longer improved model fitness, or when the parameters to search across were exhausted.

Each model was fit separately for each subject 20 times with randomized starting positions. The set of parameters with the lowest negative log-likelihood from each set of fits was saved and the other parameter fits were discarded. These 20 separate fits helped guard against the influence of idiosyncratic differences in starting point on the recovered parameters, and the possibility that the model fitting process would converge on local minima in the negative log-likelihood function.

### Parameter recovery test

To verify that all parameters of the best fitting model could be recovered, we generated 100 simulated datasets with randomized parameters. The parameter *d* was generated from a uniform distribution from 0 to 0.05. Parameters *β*_*e*_ and *β*_*s*_were generated from a lognormal distribution with a mean of 5 and standard deviation of 0.5. Parameters *β*_0_, *β*_*m*_ and *β*_*sim*_ were all generated from a standard normal distribution (mean 0, standard deviation of 1). The *c* parameter was generated from a uniform distribution ranging from 2 to 58. Given that these parameters were randomly generated and did not share the covariance of the real parameters fit to participants’ behavior, this test could overestimate their recoverability. Thus, we further verified the recoverability of these parameters by fitting the model to data simulated using the parameters fit to participants’ real behavior.

### fMRI data acquisition and preprocessing

Whole-brain imaging was acquired using a Siemens 3T Magnetom Prisma system with a 64-channel head coil. First, a high resolution T1 weighted MPRAGE image was acquired for visualization (repetition time (TR), 1900 ms; echo time (TE), 3.02 ms; flip angle, 9°; 160 sagittal slices; 1 x 1 x 1 mm voxels). Functional volumes were acquired using a gradient-echo echo planar sequence (TR, 2000 ms; TE, 28 ms; flip angle 90°; 38 interleaved axial slices tilted approximately 15° from the AC-PC plane; 3 x 3 x 3 mm voxels). Functional data were acquired over three runs. Each run lasted 16.5 min on average (Mean: 494.8 acquisitions). After the functional runs, a brief in-plane anatomical T1 image was collected, which was used to define a brain mask that respected the contours of the brain and the space of the functional runs (TR, 350 ms; TE 2.5 ms; flip angle 70°; 38 axial slices; 1.5 x 1.5 x 3 mm voxels). Soft padding was used to restrict head motion throughout the experiment. Stimuli were presented on a 32-inch monitor at the back of the bore of the magnet, and subjects viewed the screen through a mirror attached to the head coil.

Functional data were preprocessed using SPM12. Quality assurance for the functional data of each subject was first assessed through visual inspection and TSdiffAna (https://sourceforge.net/projects/spmtools/) and ArtRepair (http://cibsr.stanford.edu/tools/human-brain-project/artrepair-software.html). Outlier volumes (defined by standard deviations from the global average signal) were interpolated after preprocessing. Slice timing correction was carried out by resampling all slices to match the first slice. Next, motion during functional runs was corrected by registering volumes to the mean using rigid-body transformation. In any case where movement across runs was in excess of the voxel size in any direction (3 mm), realignment was recalculated run-by-run. Any participants where motion within a run was larger than a voxel were excluded from further analysis (N = 2). Functional volumes were then normalized to Montreal Neurological Institute (MNI) stereotaxic space using 4^th^ order B-spline interpolation, and resampled to 2 x 2 x 2 mm voxels using trilinear interpolation. Finally, volumes were spatially smoothed using an 8 mm full-width half-maximum isotropic Gaussian kernel. The in-plane anatomical T1 image for each subject was normalized to MNI space and used to create a brain mask for functional analysis using the Brain Extraction Tool in FSL (https://fsl.fmrib.ox.ac.uk/fsl/fslwiki/).

### fMRI data analysis

Functional data were analyzed under the assumptions of the GLM using SPM12. All regressors and parametric modulators were convolved with the SPM canonical hemodynamic response function (HRF). Subject motion (six translational and rotational components) were also included as nuisance regressors. Beta coefficients for single subject effects were estimated using a fixed-effects model in a first-level analysis. For whole-brain contrasts, these estimates were then included in a second-level analysis using a one-sample t-test against 0 at each voxel. Whole brain t-statistic maps were thresholded at a cluster defining threshold of *P* < 0.001 uncorrected, and then evaluated for statistical significance at the cluster corrected FWE threshold of *P* < 0.05 to control for multiple comparisons. The cluster extent for this threshold varied by test and is reported with each analysis.

### fMRI general linear models

Three different general linear models (GLMs) were fit to fMRI data:

#### (1) Condition-based GLM

The condition-based GLM included separate regressors at the response and feedback phases for the following factors: clustering (recipe, independent), value (positive, negative), presentation number (1, 2, 3) and accuracy (correct, error). The duration of the boxcar for the response regressors was set to the median of subjects’ combined reaction time and slider movement durations. Trial-by-trial deviation from this median duration was included as an additional nuisance regressor as a means of controlling for the influence of RT on a trial-to-trial basis independent of the effects of condition. This GLM also included a nuisance regressor for the time the stimulus display was on screen (i.e. the response window, ISI and feedback).

#### (2) Finite impulse response GLM

For ROI-based analyses on contrasts of task conditions, a separate finite impulse response (FIR) GLM was fit to each participant. This model had the identical conditions as the condition-based GLM described immediately above, but used an FIR model with eight time points (16 s window) as a basis function for the hemodynamic response function. This allowed us to contrast activity between conditions within different ROIs without assumptions about the shape of the hemodynamic response function made for simplification in whole-brain and model-based analyses.

#### (3) Model-based value components GLM

Finally, we tested for correlates of model-based value terms and their interaction with participants’ value ratings (i.e. participant-P(yes)) using a parametric GLM. This model had three boxcar regressors: a response regressor with onsets at stimulus presentation and durations lasting to the point participants finished moving the slider, a feedback regressor with onsets and durations lasting the presentation of feedback, and a nuisance regressor for the stimulus display. There were eight parametric modulators on the response regressor: participant-P(yes), participant confidence, experienced value, schema value, unfamiliarity, interactions between participant-P(yes) and experienced value, schema value and unfamiliarity, as well as subjective value ratings for ingredients. These interactions served as a filter to focus on neural activity that responded to the influence of these model-based terms on participants’ value judgments. We examined the covariance structure of these modulators to assess their independence. The mean *R*^2^ for all pairs of parametric modulators was 0.08, while the maximum *R*^2^ was 0.56 for the correlation between experienced value and participant-P(yes). This stronger correlation was not surprising given that experienced value was a major determinant of participants’ P(yes) ratings. The feedback regressor also had two parametric modulators: reward prediction errors (RPE; calculated as the difference between participant-P(yes) and outcomes) and surprise (calculated as the absolute value of RPEs). Parametric modulators were not orthogonalized in this model.

### Region of interest analyses

Regions of interest (ROIs) were defined for testing region-specific hypotheses using unbiased criteria. Specifically, we chose ROIs within a network previously identified by our lab and others as forming a network involved in the storage and retrieval of semantic and episodically encoded information (MTC, aVLPFC, HPC; (Badre, Poldrack, Pare-Blagoev, Insler, & Wagner, 2005a; Barredo, Verstynen, & Badre, 2016; Thompson-Schill, D’Esposito, Aguirre, & Farah, 1997)), and in forming estimates of subjective and expected value (vmPFC, mPFC, striatum; (Barron, Dolan, & Behrens, 2013; Bartra, McGuire, & Kable, 2013)). These ROIs were either defined using masks from the automated anatomical labelling atlas (AAL), or using 10 mm spheres around peak coordinates from independent studies. These spheres were then masked to only include voxels that were more than 50% likely to be gray matter in SPM’s tissue probability map. The coordinates and references for these ROIs are listed in Table 1.

**Table 1.**
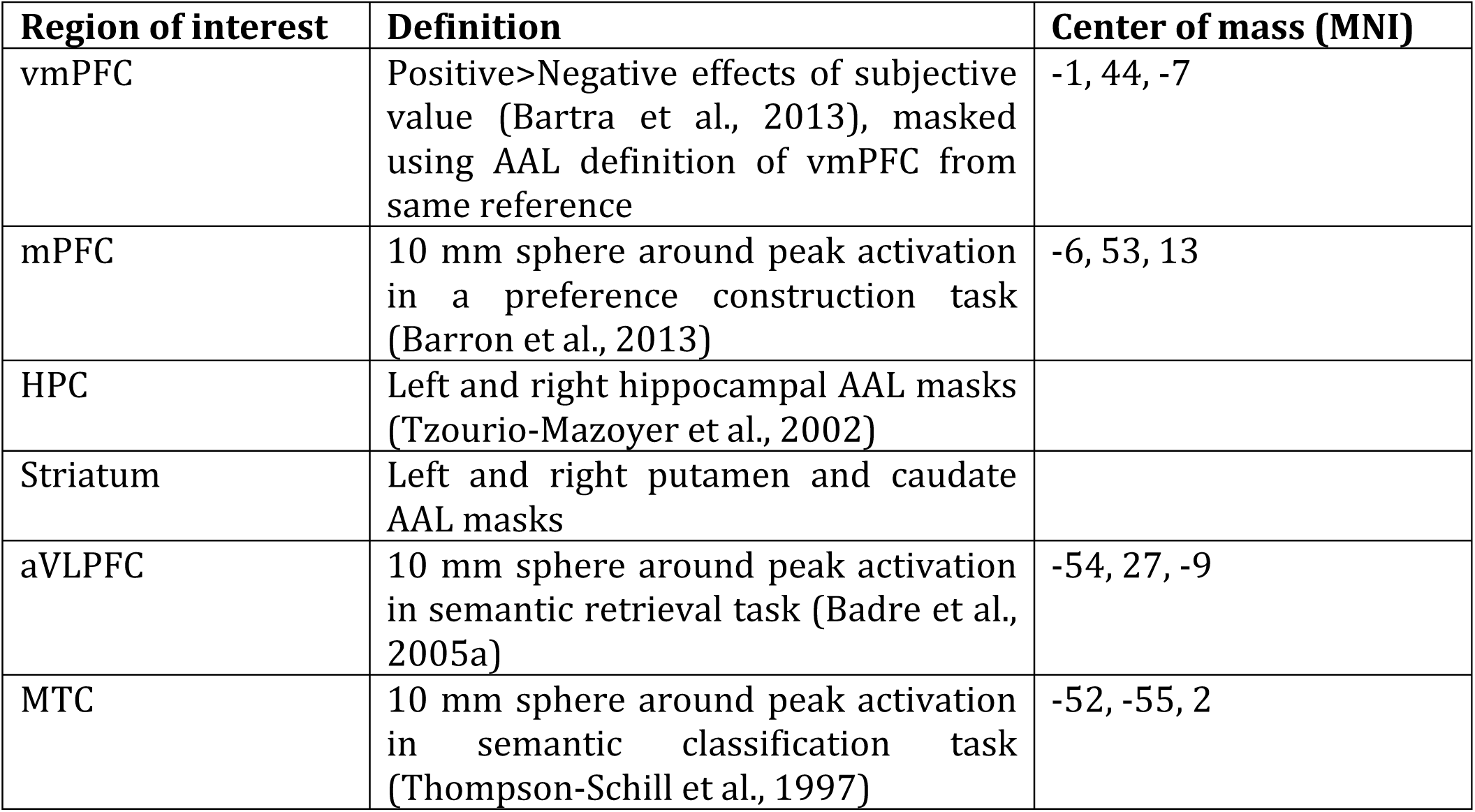
Description of ROIs

| Region of interest | Definition | Center of mass (MNI) |
| --- | --- | --- |
| vmPFC | Positive>Negative effects of subjective value (Bartra et al., 2013), masked using AAL definition of vmPFC from same reference | -1, 44, -7 |
| mPFC | 10 mm sphere around peak activation in a preference construction task (Barron et al., 2013) | -6, 53, 13 |
| HPC | Left and right hippocampal AAL masks (Tzourio-Mazoyer et al., 2002) |  |
| Striatum | Left and right putamen and caudate AAL masks |  |
| aVLPFC | 10 mm sphere around peak activation in semantic retrieval task (Badre et al., 2005a) | -54, 27, -9 |
| MTC | 10 mm sphere around peak activation in semantic classification task (Thompson-Schill et al., 1997) | -52, -55, 2 |

In ROI analyses contrasting conditions, the time course of activity within each ROI was estimated using the MarsBar Toolbox (Brett, Anton, Valabregue, & Poline, 2002) from the FIR GLM. For each ROI, the peak activity was found for using the grand-average of this timecourse across conditions and subjects (with equal weight for each conditions), and the mean activity ± 1 bin around this time point was extracted and tested using RM-ANOVAs. In ROI analyses of parametric GLM models, the mean beta weights within ROI voxels were extracted instead.

### Generalized psychophysical interaction analysis

Generalized psychophysical interaction (gPPI) analyses were carried out using the CONN toolbox in conjunction with SPM12 (McLaren, Ries, Xu, & Johnson, 2012; Whitfield-Gabrieli & Nieto-Castanon, 2012). The same ROIs described in Table 1 were used as seeds for focused ROI-ROI connectivity analyses and exploratory whole-brain analyses. Task-related activity was regressed out using the condition-based GLM described above, including six regressors for translational and rotational motion and trial-by-trial deviations from the median of reaction time and slider movement durations, as well as additional nuisance regressors including five principal components for both cerebrospinal fluid voxels and white matter, and a regressor for linear trends in the signal. The residuals of this model were then high-pass filtered at 0.008 Hz.

Following this denoising process, gPPI analyses involved a two-step process. The regressors from the condition-based GLM, the timecourse of each ROI and the interaction of this timecourse with the condition-based regressors were included in a multiple linear regression model. Task-related changes in the relationship between activity in pairs of regions was estimated using the bivariate correlation between these interactions. Correlation coefficients for each subject were Fisher transformed and contrasted between conditions. In a second level analysis, these differences were subjected to a one-sample *t*-test against zero. Exploratory seed-to-voxel whole brain connectivity analyses were similarly carried out by calculating the bivariate correlation of this interaction term between an ROI and each target voxel. These correlations were then Fisher-transformed and the difference between transformed correlations in task conditions was compared to zero with one-sample t-tests. As with above fMRI analyses, all whole-brain effects were first peak thresholded at *P* < 0.001 and further thresholded at FWE cluster corrected threshold of *P* < 0.05 to control for multiple comparisons. Cluster thresholds are reported separately for each analysis.

## Results

### Behavioral data

To test if participants took advantage of schematic knowledge to make decisions, we first examined whether participants choices reflected customers’ recipe preferences on the first presentation of ingredients before seeing any external feedback (Figure 3a). On these trials, participants accepted ingredients associated with the positive value recipe more often than ingredients associated with the negative value recipe and more often than positive value independent ingredients. This was evident in a significant three-way interaction between value, clustering and presentation number on participants’ choices (RM-ANOVA: F_2, 44_ = 27.20, *P* < 0.0001, 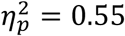), and direct contrasts between the ingredient types (Bonferroni corrected t-tests: *t*(22) ≥ 7.79, *P*’s < 0.0001, *d*’s ≥ 1.62).

**Figure 3.**
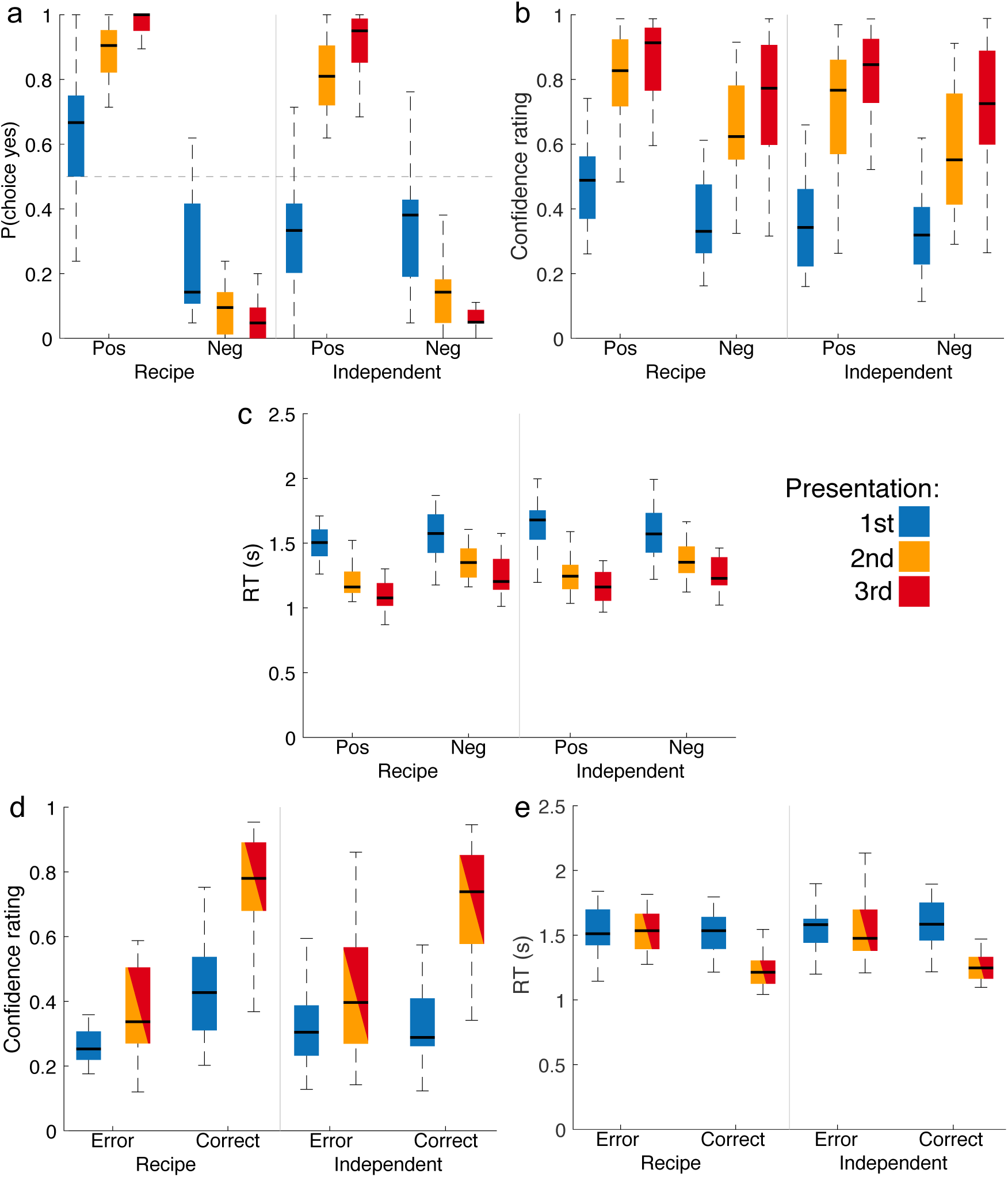
Behavioral data. Choices, reaction times (RT) and confidence ratings for positive (pos) and negative (neg) ingredients in the recipe and independent conditions across presentations. **a.** Frequency of ‘yes’ choices, **b.** Confidence ratings for correct responses. **c**. Reaction times for correct responses in recipe and independent conditions, for positive (pos) and negative (neg) ingredients. **d.** Confidence ratings for errors and correct responses in recipe and independent conditions, collapsing across value. Second and third presentations are also collapsed. **e.** Reaction times for correct and error responses in the same conditions as **d**. Boxes represent 25^th^, 50^th^ and 75^th^ quantiles of the data. Whiskers are approximately ±2.7 SD.

In contrast, there was no significant difference between acceptance rates for positive and negatively valued independent ingredients on the first presentation. This was as expected, as the ingredients in this condition did not have inferable value based on an association with a positive or negative recipe. The interaction between value and clustering on participants’ choices was confirmed with a separate RM-ANOVA on the first presentation only (F _1, 22_ = 61.31, *P* < 0.0001, 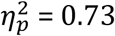). Participants accepted ingredients associated with the negative value recipe numerically less often than negative independent ingredients, but this difference was not significant after correction for multiple comparisons. Overall, the choice results indicate that participants successfully inferred value information from recipe associations, though these effects were stronger for the positive recipe.

There was clear evidence that participants learned ingredient values from feedback. Participants made choices consistent with the values of ingredients more often in the second and third presentations than in the first in all conditions, demonstrating retention of feedback from previous trials (Bonferroni corrected t-tests: *t*’s(22) ≥ 4.54, *P*’s < 0.01, *d*’s ≥ 0.95). Similarly, participants’ rate of value consistent choices increased significantly from the second presentation to the third in all conditions, except for independent positive ingredients (Bonferroni corrected t-tests: *t*’s(22) ≥ 4.69, *P*’s < 0.01, *d*’s ≥ 0.98).

Notably, participants also tended to reject both positive and negative independent ingredients in the first presentation significantly more often than chance (One sample t-tests: *t*’s(22) ≥ 8.69, *P*’s < 0.0001, *d*’s ≥ 1.81). Thus, participants strategically avoided endorsing independent ingredients before learning their values through feedback.

To test how participants’ belief in their choices changed during the task, we next examined how reported confidence changed on correct responses (Figure 3b). Confidence increased with repeated presentations, and was also greater overall for recipe and positive value ingredients (RM-ANOVA: F’s ≥ 27.88, *P* ≤ 0.0001, 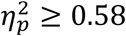). Confidence for acceptance of positive recipe ingredients was significantly higher than positive independent ingredients on the first presentation (Bonferroni corrected t-test: *t*(20) = 4.57, P < 0.01, *d* = 0.99) and third presentation (Bonferroni corrected t-test: *t*(20) = 4.96, P < 0.01, *d* = 1.08). However, there was no comparable difference in confidence for rejections of negative recipe and independent ingredients, resulting in a significant interaction between clustering and value (RM-ANOVA: F _1, 20_ = 9.28, *P* = 0.006, 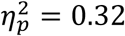),

Confidence also increased significantly from the first presentation to the second and third presentations in all conditions (Bonferroni corrected t-tests: *t*’s(20) ≥ 4.57, P’s ≤ 0.01, *d*’s ≥ 1.29), and from the second to the third presentation in all conditions, except for negative recipe ingredients (Bonferroni corrected t-tests: *t*’s(20) ≥ 3.98, P’s ≤ 0.05, *d*’s ≥ 0.87), leading to a modest interaction of value and presentation number (F _2, 40_ = 3.34, *P* = 0.04, 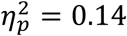).

Similar to reported confidence, participants’ reaction times were also strongly influenced by experience with ingredients with stronger decrements in subsequent presentations of positive recipe ingredients (Figure 3c). There was a significant interaction between value with presentation number (F _2, 40_ = 8.50, *P* = 0.0008, 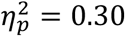), with larger reductions in reaction times for positive ingredients, consistent with better recall. There was also an interaction between value and clustering (RM-ANOVA: F _1, 20_ = 4.66, *P* = 0.04, 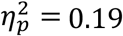), owing to comparatively bigger effect of value on recipe ingredients compared to independent ingredients. Responses in the first presentation did not significantly differ by cluster condition or value.

In anticipation of analyzing the effects of memory retrieval failures on brain activity, we also tested if errors were associated with lower confidence and longer reaction times, as would be expected for retrieval failures. We collapsed across value levels and the second and third presentations to have enough error trials per participant for this analysis. Confidence ratings were influenced by significant interactions between accuracy and clustering (Figure 3d; RM-ANOVA: F _1, 21_ = 15.43, *P* = 0.0007, 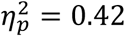), and accuracy and presentation number (F _1, 21_ = 43.69, *P* < 0.0001, 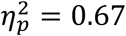). On the first presentation, participants were significantly more confident on correct responses for recipe ingredients than independent ingredients (Bonferroni corrected t-tests: *t*(21) = 3.77, P < 0.05, *d* = 0.80), and error responses on recipe ingredients (Bonferroni corrected t-tests: *t*(21) = 5.60, P < 0.01, *d* = 1.19). There was no comparable significant difference in confidence for correct and error responses for independent ingredients in the first presentation. We confirmed that the effects of accuracy differed between clustering conditions on this first presentation by separately testing this interaction (RM-ANOVA: F _1, 21_ = 14.34, *P* = 0.001, 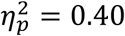). Thus, confidence on the first presentation appeared to selectively depend on retrieval of relevant recipe associations. Confidence was also higher on accurate responses compared to errors on the second and third presentations of both recipe and independent ingredients (Bonferroni corrected t-tests: *t*’s(21) ≥ 6.64, P’s < 0.0001, *d* = 1.42).

Reaction times were influenced by an interaction of accuracy and presentation number (Figure 3e; RM-ANOVA: F _1, 21_ = 49.84, *P* < 0.0001, 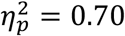), but there was only a trend toward an interaction between accuracy and clustering (F _1, 21_ = 3.81, *P* = 0.06, 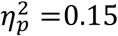), and no three-way interaction. Participants were significantly faster on correct responses than errors on the second and third presentations of both recipe and independent ingredients (Bonferroni corrected t-tests: *t*’s(21) ≥ 5.03, P’s < 0.01, *d* ≥ 1.07). This pattern is consistent with longer reaction times on errors reflecting retrieval failures (Ratcliff, 1978). In contrast, there were no significant differences in reaction times for correct and error responses in the first presentation for either clustering condition.

In summary, behavioral results established that participants were able to infer the value of ingredients based on their recipe association and used these inferred values to make choices. Participants also showed a generally risk averse attitude on this first presentation by tending to reject independent ingredients. On subsequent presentations of an ingredient, participants accurately retrieved explicit memory of deterministic feedback to make faster, more confident decisions. This was the case across both recipe and independent items. However, participants continued to benefit from recipe associations in these later presentations, with generally faster reaction times and greater confidence for correct responses in this condition compared to independent ingredients. Finally, participants tended to asymmetrically utilize knowledge about the positive recipe more than the negative recipe throughout all trials.

### Main effects of task conditions in whole-brain analyses

We hypothesized that vmPFC and striatum would respond to schema-based and experienced values of ingredients respectively, and interact with different memory systems in retrieving and constructing these values. Thus, we predicted that vmPFC activity would be modulated by schema values more than directly experienced values, and the opposite would be true of the striatum. We also predicted that a network of regions, including the striatum, HPC and middle temporal cortex would be engaged by successful retrieval (Spaniol et al., 2009), while aVLPFC would be engaged by retrieval difficulty (Badre, Poldrack, Pare-Blagoev, Insler, & Wagner, 2005b). We first tested how ingredient clustering, value, presentation number and accuracy affected activity using simple contrasts from our condition-based GLM at a whole-brain level (Table 2). These analyses were filtered to focus on correct responses, excluding trials where participants may have made errors due to failure to appropriately recall information relevant to ingredient values, except where trials with different response accuracy were directly contrasted.

**Table 2.**
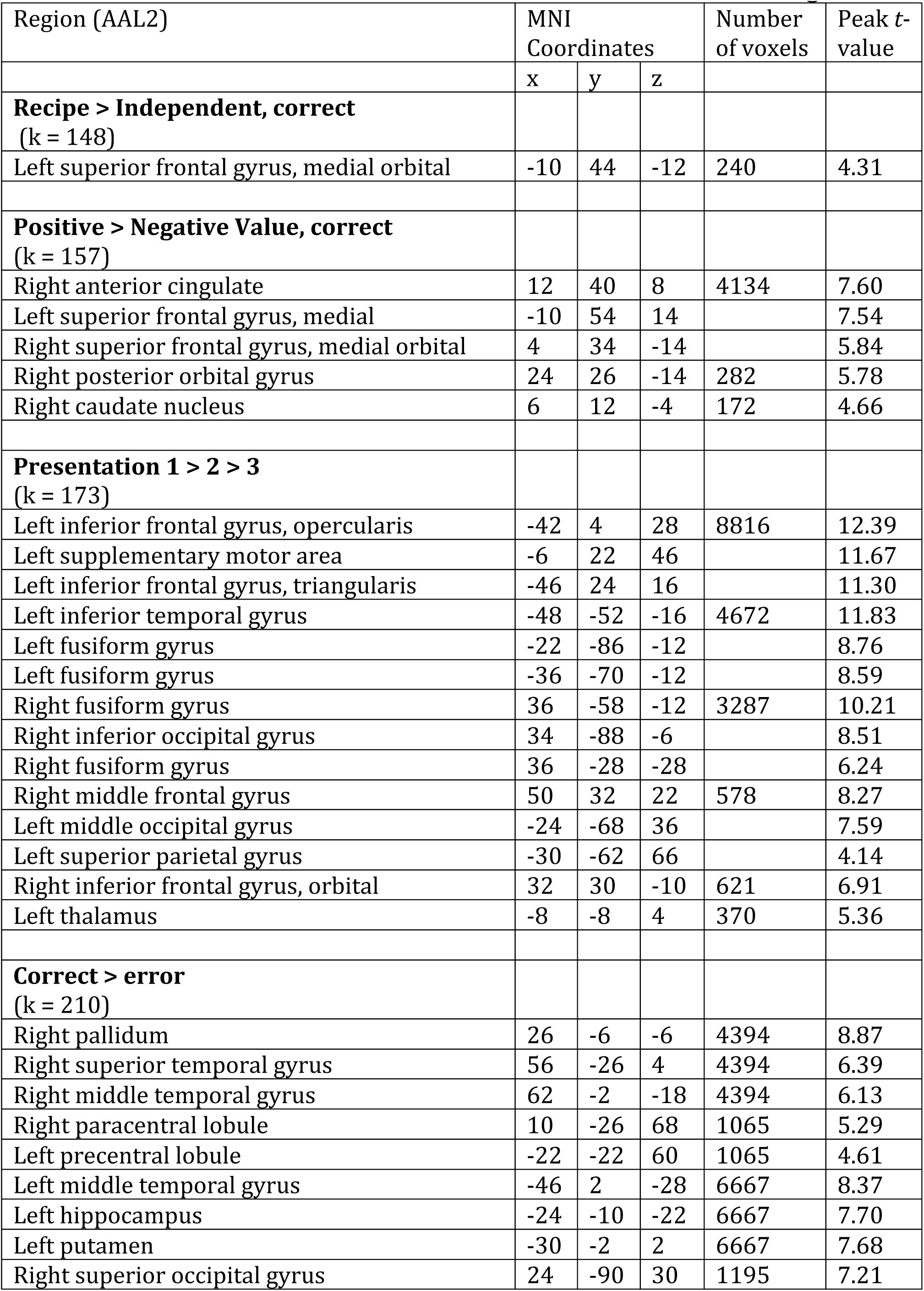
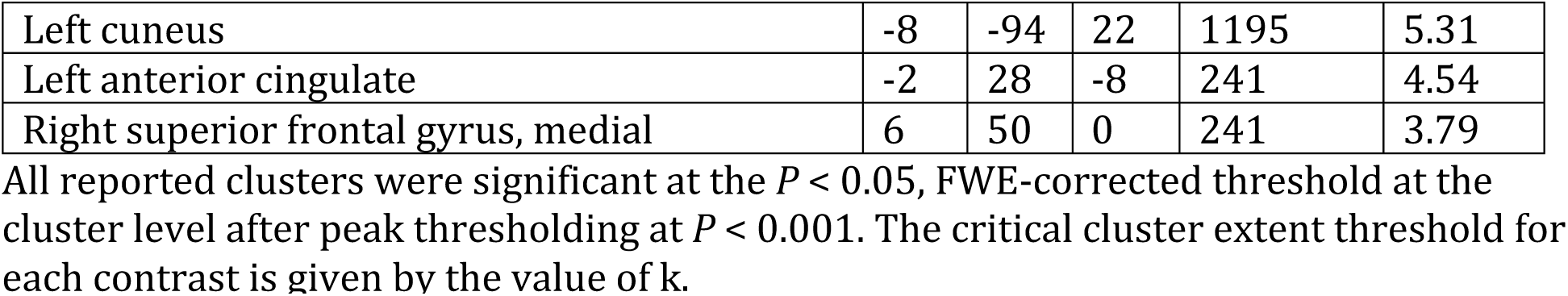
fMRI activations for main effects of task conditions shown in Figure 4

| Region (AAL2) | MNI Coordinates |  |  | Number of voxels | Peak *t*-value |
| --- | --- | --- | --- | --- | --- |
|  | x | y | z |  |  |
| **Recipe > Independent, correct**<br>(k = 148) |  |  |  |  |  |
| Left superior frontal gyrus, medial orbital | -10 | 44 | -12 | 240 | 4.31 |
| **Positive > Negative Value, correct**<br>(k = 157) |  |  |  |  |  |
| Right anterior cingulate | 12 | 40 | 8 | 4134 | 7.60 |
| Left superior frontal gyrus, medial | -10 | 54 | 14 |  | 7.54 |
| Right superior frontal gyrus, medial orbital | 4 | 34 | -14 |  | 5.84 |
| Right posterior orbital gyrus | 24 | 26 | -14 | 282 | 5.78 |
| Right caudate nucleus | 6 | 12 | -4 | 172 | 4.66 |
| **Presentation 1 > 2 > 3**<br>(k = 173) |  |  |  |  |  |
| Left inferior frontal gyrus, opercularis | -42 | 4 | 28 | 8816 | 12.39 |
| Left supplementary motor area | -6 | 22 | 46 |  | 11.67 |
| Left inferior frontal gyrus, triangularis | -46 | 24 | 16 |  | 11.30 |
| Left inferior temporal gyrus | -48 | -52 | -16 | 4672 | 11.83 |
| Left fusiform gyrus | -22 | -86 | -12 |  | 8.76 |
| Left fusiform gyrus | -36 | -70 | -12 |  | 8.59 |
| Right fusiform gyrus | 36 | -58 | -12 | 3287 | 10.21 |
| Right inferior occipital gyrus | 34 | -88 | -6 |  | 8.51 |
| Right fusiform gyrus | 36 | -28 | -28 |  | 6.24 |
| Right middle frontal gyrus | 50 | 32 | 22 | 578 | 8.27 |
| Left middle occipital gyrus | -24 | -68 | 36 |  | 7.59 |
| Left superior parietal gyrus | -30 | -62 | 66 |  | 4.14 |
| Right inferior frontal gyrus, orbital | 32 | 30 | -10 | 621 | 6.91 |
| Left thalamus | -8 | -8 | 4 | 370 | 5.36 |
| **Correct > error**<br>(k = 210) |  |  |  |  |  |
| Right pallidum | 26 | -6 | -6 | 4394 | 8.87 |
| Right superior temporal gyrus | 56 | -26 | 4 | 4394 | 6.39 |
| Right middle temporal gyrus | 62 | -2 | -18 | 4394 | 6.13 |
| Right paracentral lobule | 10 | -26 | 68 | 1065 | 5.29 |
| Left precentral lobule | -22 | -22 | 60 | 1065 | 4.61 |
| Left middle temporal gyrus | -46 | 2 | -28 | 6667 | 8.37 |
| Left hippocampus | -24 | -10 | -22 | 6667 | 7.70 |
| Left putamen | -30 | -2 | 2 | 6667 | 7.68 |
| Right superior occipital gyrus | 24 | -90 | 30 | 1195 | 7.21 |
| Left cuneus | -8 | -94 | 22 | 1195 | 5.31 |
| Left anterior cingulate | -2 | 28 | -8 | 241 | 4.54 |
| Right superior frontal gyrus, medial | 6 | 50 | 0 | 241 | 3.79 |

All reported clusters were significant at the $P < 0.05$ , FWE-corrected threshold at the cluster level after peak thresholding at $P < 0.001$ . The critical cluster extent threshold for each contrast is given by the value of $k$ .

Recipe ingredients elicited greater activity than independent ingredients in medial OFC (Figure 4a), but not elsewhere. Ingredients associated with positive value elicited greater activity in OFC and medial PFC, as well as the ventral striatum (Figure 4b). Compared with later ingredient presentations, the first encounter with an ingredient was associated with widespread activity in lateral PFC, including left VLPFC, inclusive of the aVLPFC, and the caudate nucleus (Figure 4c). Activity in left MTC, bilateral HPC and putamen was higher in correct responses compared to errors across all presentations (Figure 4d).

**Figure 4.**
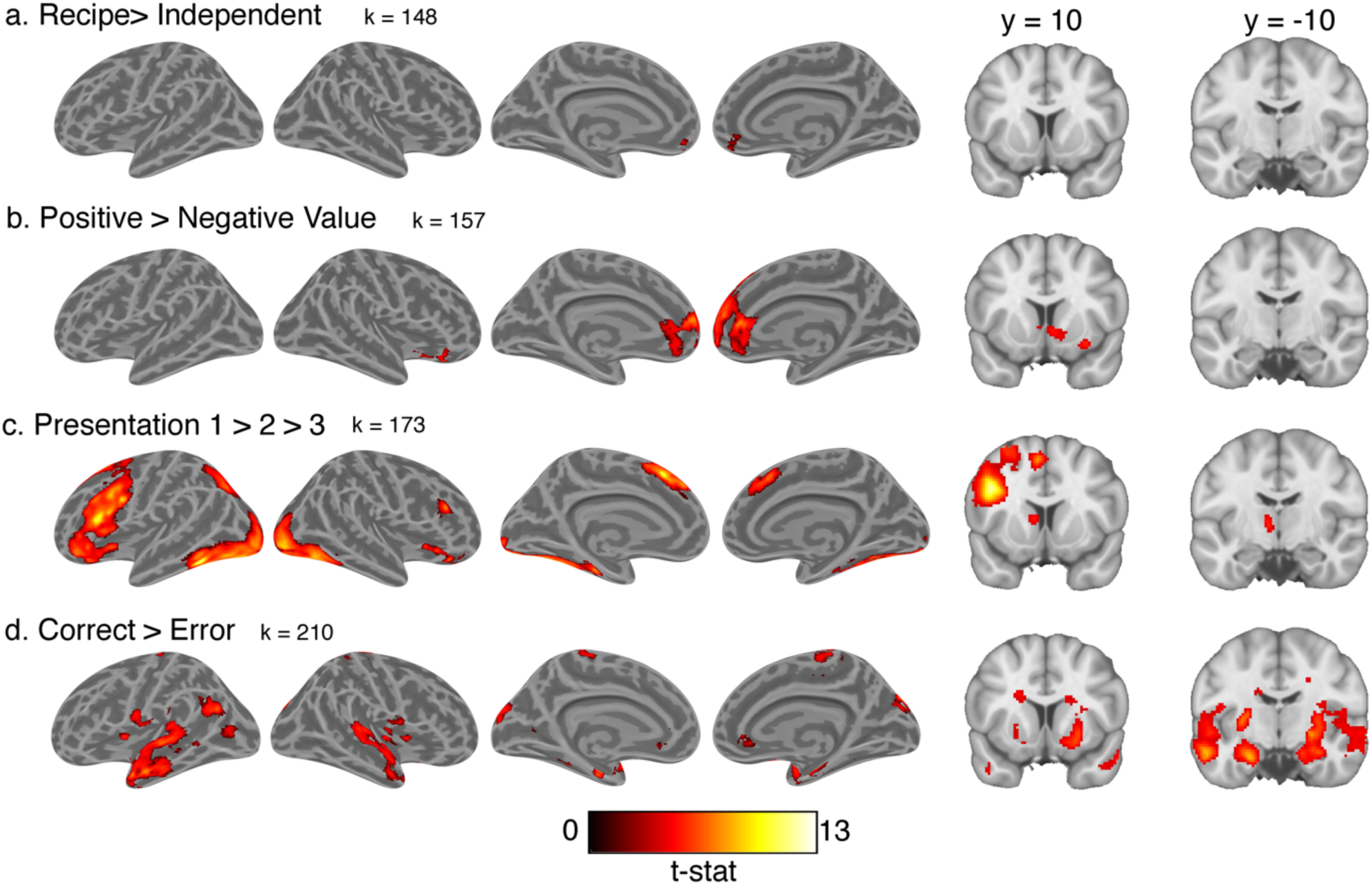
Whole brain contrasts projected onto inflated cortical surfaces and coronal slices for main effects of **a**. ingredient-recipe clustering, **b**. ingredient value, **c**. presentation number and **d**. response accuracy. All images are thresholded at *P* < 0.05, FWE cluster corrected. All main effects, except for accuracy, are filtered to only include correct responses. Cluster extent threshold for each contrast is given by value of k.

### Region of interest analyses

To follow-up the whole-brain voxel-wise contrasts, we tested for more specific task-dependent changes in activity within a set of *a priori* regions of interest (ROIs) defined in Table 1 using a finite impulse response (FIR) model of the hemodynamic response. We were primarily interested in how the clustering status of ingredients influenced value-related activity across ingredient repetitions on correct responses. Here we focused on vmPFC and the striatum, since we had *a priori* expectations that these regions would differentially process value related to episodic and schematic memory. As the medial frontal wall is functionally heterogeneous (Braga & Buckner, 2017; Roy, Shohamy, & Wager, 2012), we included two ROIs from this region, one that was defined by a meta-analysis of studies of expected-value (Bartra et al., 2013), and a more dorsal ROI (mPFC) that responded to shared associations between components of a recipe schema in another experiment and was independent and non-overlapping with the vmPFC ROI (Barron et al., 2013). This dorsal mPFC ROI responded to ingredient value, but not any other task manipulations.

vmPFC was more active for positive than negative expected value, evident in a main effect of value (Figure 5a; RM-ANOVA: F _1, 20_ = 9.13, *P* = 0.007, 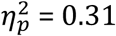), consistent with the whole-brain analysis. Activity in this region was greater for positive compared to negative recipe ingredients in the third presentation (Bonferroni corrected t-test: *t*(20) = 4.90, P < 0.01, *d* = 1.07), but there was no comparable value difference for independent ingredients (Bonferroni corrected t-test: *t*(20) = 1.13, P > 0.05, *d* = 0.25). There was no significant difference between vmPFC activity in response to positive and negative recipe ingredients on the first presentation, leading to a three-way interaction between value, presentation number and clustering (F _1, 20_ = 5.06, *P* = 0.01, 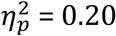). This null result of value on the first presentation was not merely due to the ROI selection; an exploratory whole-brain contrast of positive and negative recipe conditions on the first presentation in the condition-based GLM was similarly null. vmPFC activity was also generally higher in later presentations, reflected in a marginal main effect of presentation number (F _2, 20_ = 3.29, *P* = 0.05, 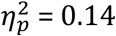), but there was no significant main effect of clustering (F _1, 20_ = 1.42, *P* = 0.2, 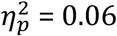).

**Figure 5.**
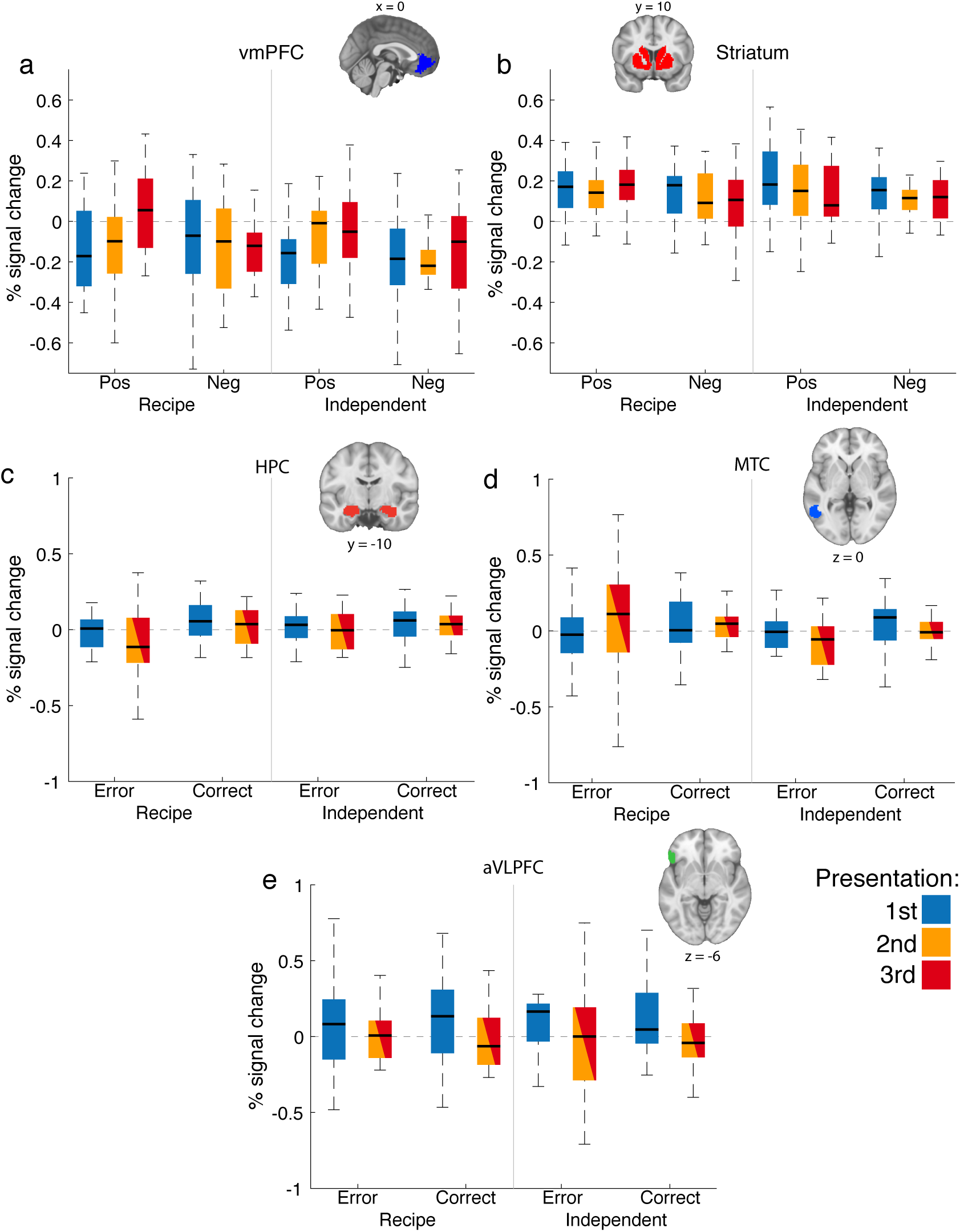
Region-of-interest analyses showing percent signal change in **a.** ventromedial prefrontal cortex (vmPFC) and **b.** the bilateral striatum on correct response trials for positive (pos) and negative (neg) ingredients in the recipe and independent conditions across presentations. **c.** Percent signal change in the hippocampus (HPC) for correct and erroneous responses in the recipe and independent conditions, collapsed across value conditions. Second and third presentations are collapsed. **d.** Middle temporal cortex (MTC) and **e.** anterior ventrolateral prefrontal cortex (aVLPFC), same conditions as **c**.

Within the striatum ROI, there was greater activation for positive than negative value, evident in a strong main effect of value (Figure 5b; F _1, 20_ = 9.17, *P* = 0.007, 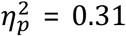), consistent with the observation from the whole brain voxel-wise analysis. However, there were no other significant main effects or interactions.

We hypothesized that value judgments in this task would rely on different neural systems supporting different types of memory. In particular, we predicted that retrieval of values associated with independent ingredients learned through feedback would place greater demands on the HPC, while the MTC would be involved more in trials where semantic memory about recipes was needed to infer ingredient values and aVLPFC would be engaged by greater retrieval demands on the first presentation. To test these predictions, we evaluated the effects of clustering status, presentation number and accuracy in ROI-based analyses of the HPC, MTC and aVLPFC. We collapsed across value conditions and combined presentations two and three into a single level to have enough error trials for this analysis.

HPC activity was greater on accurate trials, confirming the whole-brain results (Figure 5c; RM-ANOVA: F _1, 21_ = 13.14, *P* = 0.002, 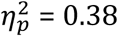). HPC activity was also generally lower on later presentations (F _1, 21_ = 6.54, *P* = 0.02, 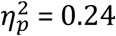). The effect of accuracy on HPC activity was modulated marginally by clustering (F _1, 21_ = 4.65, *P* = 0.04, 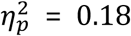). Surprisingly, we found greater HPC activity on correct trials than error trials for recipe ingredients, collapsing across presentations (Bonferroni corrected t-test: *t*(21) = 3.47, P < 0.01, *d* = 0.74), but no such accuracy effect for independent ingredients. There was no main effect of clustering, or three-way interaction of clustering, accuracy and presentation.

MTC activity responded to an interaction of clustering and presentation number (Figure 5d; F _1, 21_ = 5.60, *P* = 0.03, 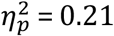). Posthoc tests collapsed across accuracy revealed a significantly greater reduction in MTC activity for independent ingredients in the second and third presentations compared to the first presentation (Bonferroni corrected t-test: *t*(21) = 2.87, P < 0.05, *d* = 0.61), but no significant difference for recipe ingredients. While MTC seemed to be engaged on the first presentation in all conditions, this reduced in subsequent presentations for independent ingredients, consistent with less reliance on semantic knowledge in this condition.

aVLPFC activity was greater on the first presentation compared to the second and third, consistent with the whole-brain analysis (Figure 5e; Presentation Main Effect: F _1, 21_ = 10.76, *P* = 0.004, 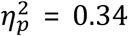). The response of aVLPFC did not change with accuracy or clustering status, or with any interaction between task variables.

In summary, ROI-based analyses confirmed whole-brain contrasts demonstrating that the vmPFC and striatum carried information about ingredient values. Unlike the striatum, value-related activity in vmPFC was greater for recipe ingredients compared to independent ingredients in later presentations, but not in the first presentation where the demand for inferring ingredient values from recipe associations was the greatest. In line with our predictions, the MTC responded to differential demands on semantic memory, though this was only true in later presentations. aVLPFC activity diminished with presentation overall, perhaps in line with greater retrieval demands early on. While the HPC responded differentially to retrieval success and failures in later presentations, this effect was surprisingly greater for recipe ingredients than independent ingredients, possibly due to the stronger encoding of these recipe ingredients, evident in participants’ behavior on these trials.

### Computational model comparison

We had hypothesized that vmPFC would be preferentially engaged by schema-based values, while the striatum would be involved in evaluating the contents of episodically encoded value information. While value-related differences in vmPFC activity grew larger in later presentations of recipe ingredients compared to independent ingredients, this change in activity could not be easily differentiated from value information learned from feedback. Importantly, this condition-based analysis also ignores the continuous nature of associations between independent ingredients and recipes. For example, strawberries are considered part of the ‘cake’ cluster, but also have some association with ‘pie.’ Further, filtering trials by accuracy based on participant responses and task-defined values ignores that responses to independent ingredients on the first presentation are made without any useful experienced or schematic value information, and ignores systematic biases or strategies participants may use in making choices that are not explicitly captured by our experimental design (e.g. a bias against endorsing independent ingredients on their first presentation).

To better separate these components of expected value for testing our first two hypotheses regarding the neural systems supporting valuation of information from different memory sources, we developed a computational model of behavior to estimate the contribution of information learned directly from experience and inferred from schematic associations to participants’ choices in each trial. We then used these values in a model-based parametric GLM to determine the correlates of this information during value judgment.

Our computational model of behavior made a few basic assumptions about learning in this task that are described in more detail in the Materials and Methods section. Briefly, each ingredient had an ‘experienced value’ (*EV*), which depended on feedback and decayed over time, and a ‘schema value’ (*SV*), which was derived from the differences in associative strength between an ingredient and the positive and negative recipes. The associative strength of an ingredient with each recipe was based on a latent semantic analysis of normative ingredient-recipe associations (Dumais, 2004; Landauer et al., 1998). Each of these value terms were weighted by a temperature parameter that determined their contribution to the model’s output, a continuous estimate of the probability that it would endorse an ingredient, which we will refer to as model-P(yes).

Starting with these assumptions, we followed a step-wise process of adding parameters to a model with these features and using cross-validation to compare the new model against the best simpler model. This allowed us to test if the new model better explained participants’ behavior on trials that were held out from the dataset used to fit parameter values. Figure 6 shows the difference in the negative log likelihoods for these alternative models against the base model. Please see the methods section for a fuller accounting of the parameters in the best fitting model and alternative models in this figure.

**Figure 6.**
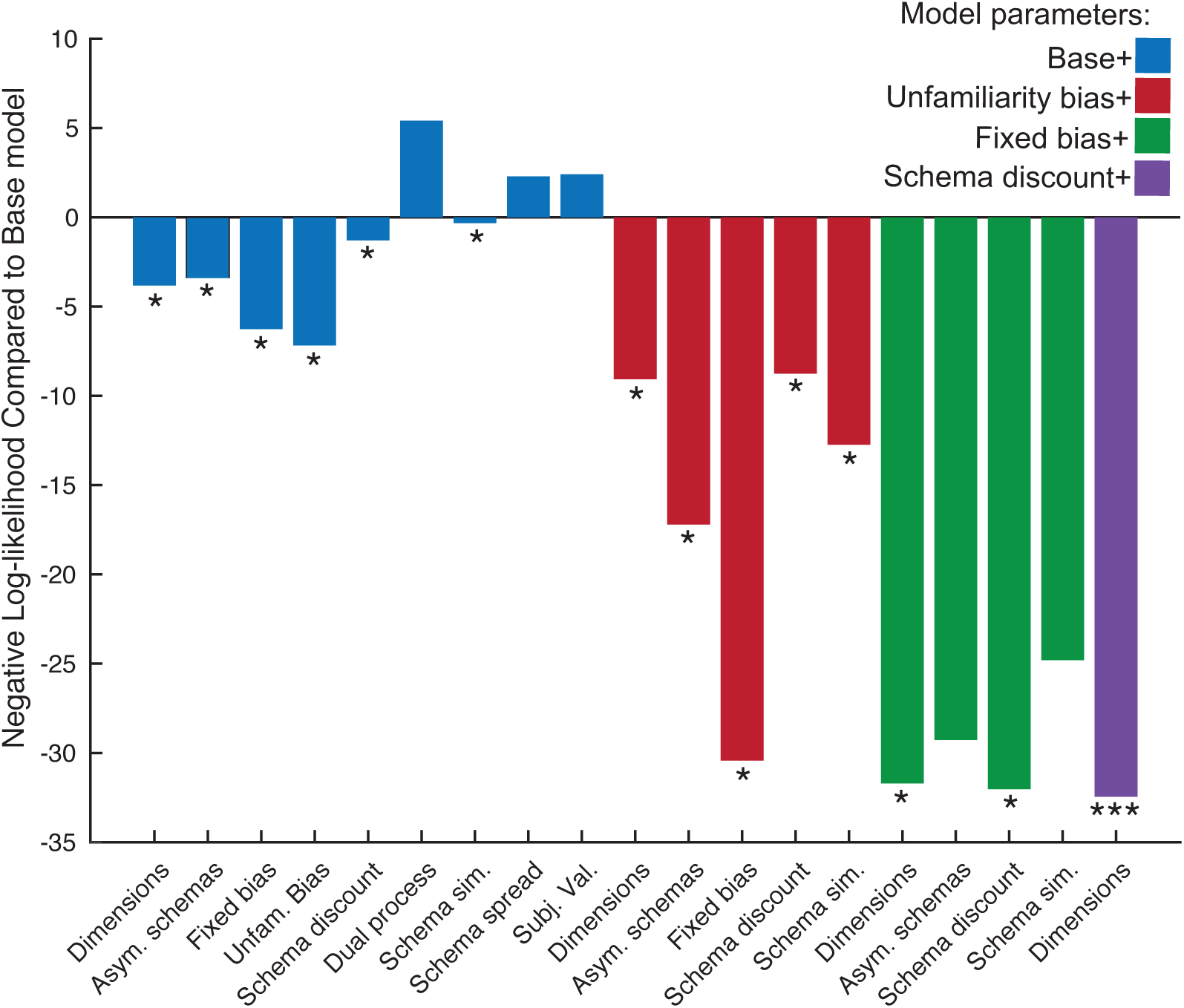
Difference in negative log-likelihood for held-out data in cross-validation procedure for additional model parameters compared to base model. Model parameters were added in a step-wise process where the parameters that best improved on the previous best fitting model were added to the next step (indicated in the legend). See Materials and Methods section for a more complete description of these models. * Reduction in negative log-likelihood compared to next best fitting model. ***, best model within a step and an improvement in fitness compared to next best model.

The final best fitting model allowed for a more sophisticated parameterization of SV where the number of dimensions in the latent semantic analysis of recipe-ingredient associations was allowed to vary. This parameter essentially controlled the depth of recipe information considered in computing *SV* (from simple distinctions like savory-sweet to more fine-grained information about particular recipes; Figure 2). The model also discounted the influence of *SV* based on the memory strength of the ingredients’ experienced value, with schema values contributing less to ingredient values when these experienced values were stronger.

*Unfamiliarity* with an ingredient was also allowed to influence decisions about betting or passing in the best fitting model. Unfamiliarity was operationalized as the opposite of the unsigned experienced value of items. In other words, unfamiliarity was highest when ingredients were seen on the first presentation, or when ingredients had not been seen for many trials and memory for their experienced value had decayed. This parameter thus effectively accounted for variance in participants’ willingness to bet on ingredients when information about their experienced values was weaker. Lastly, an additional free parameter acted as an intercept term in the decision function, which effectively acted as a fixed bias on the model’s tendency to bet on ingredients.

### Computational model validation

To ensure that our computational model was capturing participants’ decisions and understand how it was accomplishing this, we carried out a series of analyses to validate its parameters and behavior. We first examined if there were general tendencies for the two bias and schema similarity temperature parameters towards positive or negative values by conducting two-tailed, one-sample t-tests against zero (Table 3). While the unfamiliarity bias parameter was significantly lower than zero (*t*(22) = 4.13, *P* = 0.0004, *d* = 0.86), the opposite was true of the fixed bias parameter (*t*(22) = 4.69, *P* = 0.0001, *d* = 0.98).

**Table 3.** Mean parameter values and Pearson correlations between parameters in the best-fitting behavioral model

|  | Mean<br>(SD) | Decay | EV<br>temp. | SV<br>temp. | UF temp. | Fixed<br>bias | Schema<br>discount<br>(log-space) |
| --- | --- | --- | --- | --- | --- | --- | --- |
| **Decay** | **0.02**<br>**(0.01)** | - |  |  |  |  |  |
| **EV temp.** | **5.18**<br>**(0.31)** | 0.28 | - |  |  |  |  |
| **SV temp.** | **2.45**<br>**(1.70)** | 0.04 | 0.37 | - |  |  |  |
| **UF temp.** | **-0.84**<br>**(0.97)** | 0.12 | 0.27 | 0.21 | - |  |  |
| **Fixed bias** | **0.69**<br>**(0.70)** | 0.06 | -0.29 | -0.29 | -0.94*** | - |  |
| **Schema**<br>**discount**<br>**(log-space)** | **-2.47**<br>**(6.05)** | -0.20 | 0.28 | 0.15 | -0.03 | -0.06 | - |
| **Recipe**<br>**dimensions**<br>**(log-space)** | **1.10**<br>**(0.36)** | -0.12 | -0.05 | 0.40 | -0.37 | 0.24 | 0.36 |

Experienced value (EV), schema value (SV), unfamiliarity (UF), \*\*\* $P < 0.0001$

It is possible that this unfamiliarity term simply captures a bias to pass on the first presentation of ingredients rather than an effect of unfamiliarity more generally — i.e. the additional effect of the passage of time and repeated presentations. To examine if there was an effect of unfamiliarity on participants’ willingness to endorse an ingredient in later trials, we used a logistic regression in each participant to test if there was a relationship between the model-based unfamiliarity term on betting decisions on the second and third presentations. A one-sample t-test revealed that these logistic regression coefficients were significantly lower than zero across participants, indicating a persistent negative influence for unfamiliarity on participants’ decisions in later presentations (*t*(22) = 3.62, P = 0.001, *d* = 0.75). Thus, this unfamiliarity term captures additional variance in decision behavior that beyond an aversion to betting on the first presentation of an ingredient.

We also examined the covariance between parameters recovered from fitting this model to participants’ behavioral data to test their independence (Table 3). The fixed bias parameter had a strong negative correlation with the unfamiliarity bias parameter, indicating that this fixed bias in effect acted to counter the aversion to betting related to unfamiliarity. Despite this strong relationship, both parameters resulted in large improvements in model fitness when included together (Figure 6), indicating that they were complimentary.

To confirm that all of these parameters could be uniquely recovered in our model fitting procedure, we generated 100 simulated participants with randomly assigned parameters using the best fitting model. We reliably and accurately recovered all parameters in these data, with high correlations between simulated and recovered parameters (Pearson *r*’s(98) ≥ 0.82, *P*’s < 0.0001). However, as these data were simulated from randomly generated parameters, they did not share the covariance structure of the parameter estimates fit to participants’ behavior described above. To assure that we could recover these parameters with the same covariance, we fit the model to simulated data generated using parameter estimates from the model fit to participants’ behavior. For all seven parameters, there was a near perfect correlation between the parameter estimates from participants’ behavior and the simulated data (Pearson *r*’s(21) ≥ 0.99, *P*’s < 0.0001).

Finally, we generated simulated behavior from the model using participants’ recovered parameters to test that the model recapitulated key features of participants’ behavior across task conditions. To directly compare the model and participants’ behavior, we focused on participants’ continuous estimate of whether or not they should endorse an ingredient, based on position of the slider between the ‘yes’ and ‘no’ ends of the bar (i.e. participant-P(yes)). We compared these ratings to the model-P(yes) measure, and transformed model-P(yes) to confidence based on the distance from the point of maximum uncertainty (i.e. 0.5). Participant-P(yes) ratings followed a similar pattern to their averaged binary choices, including key features on the first presentation like higher P(yes) for positive recipe ingredients compared to negative recipe, and positive independent, ingredients, and a bias against endorsing independent ingredients (data not shown)

We analyzed model-P(yes) and confidence ratings in the same way as participants’ responses. Like with participants’ choices, there was a strong three-way interaction between value, clustering and presentation number (Figure 7a; RM-ANOVA: F _2, 44_ = 73.09, *P* < 0.0001, 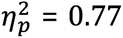). The model used recipe information to make decisions on the first presentation, with higher model-P(yes) ratings for positive recipe ingredients over negative recipe ingredients, and over independent positive ingredients (Bonferroni corrected t-tests: *t*’s(22) ≥ 9.12, P’s < 0.0001, *d*’s ≥ 1.90). There was also no significant difference in model-P(yes) for positive and negative independent ingredients on this first presentation. Unlike real participants, the model gave significantly lower ratings to negative recipe ingredients than negative independent ingredients on the first presentation (Bonferroni corrected t-test: *t*(22) = 9.52, P < 0.0001, *d* = 1.99), suggesting that it used information about the negative recipe more than participants did. Model-P(yes) on the first presentation of independent ingredients trended negatively, but was not significantly lower than chance (*t*(22) = 1.65, P = 0.1, *d* = 0.34), indicating a modest difference from participant behavior. In subsequent presentations, the model also followed the values of options learned through feedback, hewing closer to ingredient values in presentation two and three compared to one (Bonferroni corrected t-tests: *t*’s(22) ≥ 8.19, P’s < 0.0001, *d* ≥ 1.71), and presentation three compared to two (Bonferroni corrected t-tests: *t*’s(22) ≥ 6.03, P’s < 0.01, *d* ≥ 1.26).

**Figure 7.**
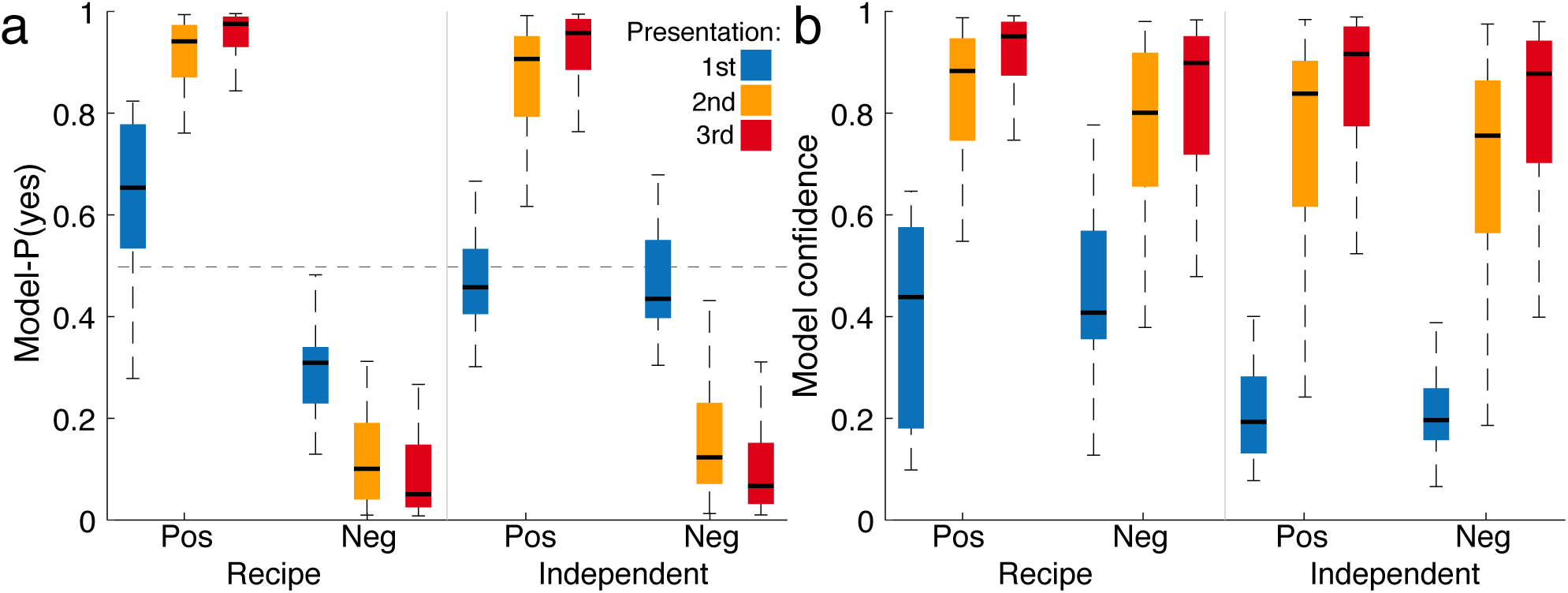
Model behavior for positive (pos) and negative (neg) ingredients in the recipe and independent conditions across presentations. **a.** Model-P(yes), a continuous indicator of the model’s estimate of the value of these options. **B.** Model confidence on correct responses only.

Similarly, we examined whether the model’s predicted confidence followed the same pattern as participants’ responses across task conditions on trials where participants responded correctly (Figure 7b). Model confidence rose predictably with presentation number (RM-ANOVA: F _2, 40_ = 165.64, *P* < 0.0001, 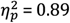), and was higher overall for recipe ingredients than independent ingredients (F _1, 20_ = 46.48, *P* < 0.0001, 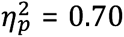), however there was no main effect of value (F _1, 20_ = 1.96, *P* = 0.2, 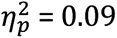). Model confidence was also notably higher for both positive and negative recipe ingredients compared to independent ingredients on the first presentation (Bonferroni corrected t-tests: *t*’s(20) ≥ 4.50, P’s < 0.01, *d*’s ≥ 0.98), rather than just positive recipe ingredients.

In summary, our model captured most core aspects of participants’ behavior in this task, including utilization of recipe-based values. The model was also able to recreate the basic pattern of participants’ reported confidence rising with repeated presentations, with benefits for positive value and recipe associations. However, the model gave greater weight to the negative value recipe than participants and underestimated the effects of value on confidence. While an alternative model with asymmetric weights for positive and negative recipes improved model fitness, this addition did not survive our model selection process (Figure 6), possibly due to variance between participants in the extent of this asymmetry. The model also underestimated the effects of unfamiliarity on P(yes) on the first presentation of independent ingredients, though was trending in the same direction as participants.

### Model-based fMRI

We had hypothesized that vmPFC would be engaged more by values inferred from schematic associations than directly experienced values, which we hypothesized would be tracked by activity in the striatum. Our computational model showed that ingredient values could be captured by a combination of experienced value, schema value and unfamiliarity, and allowed us to leverage these terms in testing these hypotheses by examining their correlates in brain activity.

To focus on brain activity that responded to the influence of these model-based value terms on participants’ value judgments, we included interactions between these three terms and participant-P(yes) in our GLM. These interactions selectively captured activity that responded to the conjunction (or agreement) between the participants’ responses with experienced and schema values (e.g. when schema value was high and participant-P(yes) were high, or when both were low). The interaction of unfamiliarity with value ratings similarly let us capture where activity increased in response to higher participant-P(yes) when unfamiliarity was high, or lower participant-P(yes) when unfamiliarity was low. In doing so, we could focus on activity where behavior and model predictions were in agreement. In contrast, the main effects of these model-based terms only captured the extent to which correlates of this information were present in the brain, independent of participants’ response. This analysis may seem circular, as we are looking for correlates of an interaction between these model-based terms with the behavior to which these terms were fit. However, the fMRI correlates that are the dependent measure in this analysis were not part of the model fitting, breaking any potential circularity in our results. This GLM also included parametric modulators for participants’ confidence ratings and participants’ subjective value ratings (provided separately in a different task) on the response regressor as nuisance terms.

Whole-brain cluster corrected analyses found no significant main effects of participant-P(yes), experienced or schema-based value, or participants’ subjective value ratings. There were significant correlates for the main effect of unfamiliarity, with increased activity in lateral PFC, visual cortex and several other regions, similar to the main effect of presentation number in the condition-based contrast above. There were also significant main effects for response confidence in the superior frontal gyrus and visual cortex (Table 4).

**Table 4.** fMRI activations for model-based values GLM, including those from Figure 8

| Region (AAL2) | MNI Coordinates |  |  | Number of voxels | Peak *t*-value |
| --- | --- | --- | --- | --- | --- |
|  | x | y | z |  |  |
| **Unfamiliarity**<br>(k = 161) |  |  |  |  |  |
| Left fusiform gyrus | -36 | -44 | -20 | 5443 | 11.02 |
| Left inferior occipital gyrus | -28 | -84 | -10 | 5443 | 9.20 |
| Left middle temporal gyrus | -60 | -48 | -8 | 5443 | 7.03 |
| Right fusiform gyrus | 34 | -52 | -12 | 3877 | 9.98 |
| Right lingual gyrus | 24 | -88 | -6 | 3877 | 9.50 |
| Right fusiform gyrus | 34 | -28 | -28 | 3877 | 7.42 |
| Left inferior frontal gyrus, orbitalis | -40 | 32 | -12 | 2995 | 9.54 |
| Left inferior frontal gyrus, opercularis | -40 | 4 | 28 | 2995 | 9.17 |
| Left inferior frontal gyrus, triangularis | -42 | 24 | 14 | 2995 | 8.45 |
| Left superior frontal gyrus, dorsolateral | -16 | 38 | 50 | 1843 | 7.67 |
| Left middle frontal gyrus | -32 | 18 | 58 | 1843 | 5.32 |
| Left supplementary motor area | -6 | 12 | 56 | 1843 | 7.15 |
| Left gyrus rectus | -2 | 48 | -18 | 403 | 7.08 |
| Left precuneus | -4 | -54 | 18 | 173 | 6.80 |
| Left hippocampus | -18 | -8 | -20 | 173 | 6.31 |
| Right gyrus rectus | 2 | 28 | -20 | 173 | 3.94 |
| **Confidence**<br>(k = 158) |  |  |  |  |  |
| Left superior frontal gyrus, medial | -6 | 50 | 46 | 240 | 5.84 |
| Left superior frontal gyrus, medial | -4 | 56 | 22 | 240 | 4.01 |
| Left middle temporal gyrus | -58 | -12 | -16 | 187 | 4.25 |
| Left middle temporal gyrus | -64 | -30 | 0 | 159 | 4.87 |
| Right Postcentral gyrus | 24 | -32 | 64 | 196 | 4.58 |
| **Schema value x participant-P(yes)**<br>(k = 161) |  |  |  |  |  |
| Right middle temporal gyrus | 52 | -42 | 4 | 559 | 5.789 |
| Right inferior temporal gyrus | 58 | -60 | -14 | 559 | 4.243 |
| Right inferior frontal gyrus | 48 | 36 | 0 | 229 | 5.664 |
| Left superior temporal gyrus | -54 | -6 | -2 | 256 | 5.168 |
| Left middle temporal gyrus | -58 | 4 | -22 | 256 | 3.93 |
| Right superior temporal gyrus | 66 | -10 | 2 | 269 | 5.015 |
| Left cuneus | 2 | -82 | 38 | 301 | 4.872 |
| Right cuneus | 18 | -86 | 26 | 301 | 3.882 |
| Left middle temporal gyrus | -66 | -50 | 2 | 250 | 4.747 |
| Right supramarginal gyrus | 58 | -26 | 20 | 182 | 4.682 |
| Left superior temporal gyrus | -64 | -30 | 18 | 193 | 4.657 |
| **Experienced value x participant-**<br>**P(yes)**<br>**(k = 168)** |  |  |  |  |  |
| Left superior frontal gyrus, medial | -4 | 60 | 8 | 1198 | 6.92 |
| Left gyrus rectus | -8 | 38 | -16 | 1198 | 4.95 |
| Right superior frontal gyrus, medial | 10 | 58 | 28 | 1198 | 4.18 |
| Left angular gyrus | -50 | -76 | 24 | 185 | 5.21 |

All reported clusters were significant at the $P < 0.05$ , FWE-corrected threshold at the cluster level after peak thresholding at $P < 0.001$ . The critical cluster extent threshold for each contrast is given by the value of k.

The interaction of schema value and participant-P(yes) was associated with activity in the left and right middle temporal gyrus, right VLPFC and bilateral supramarginal gyri (Figure 8a). The interaction of experienced value and participant P(yes) was also associated with activity in medial OFC and a large area of the medial superior frontal gyrus and left angular gyrus (Figure 8b). There were no clusters that passed the whole-brain FWE cluster-corrected threshold for the interaction of unfamiliarity and participant-P(yes). However, given the challenge of detecting cluster corrected effects in deep brain structures, we applied a small volume correction using the striatum ROI. This analysis found that a cluster in the bilateral caudate nucleus responded to the interaction of unfamiliarity with participant-P(yes) (Figure 8c; FWE-corrected *P* < 0.05, MNI: 18, 20, −4). Whole brain peak activations for this GLM are described in Table 4.

**Figure 8.**
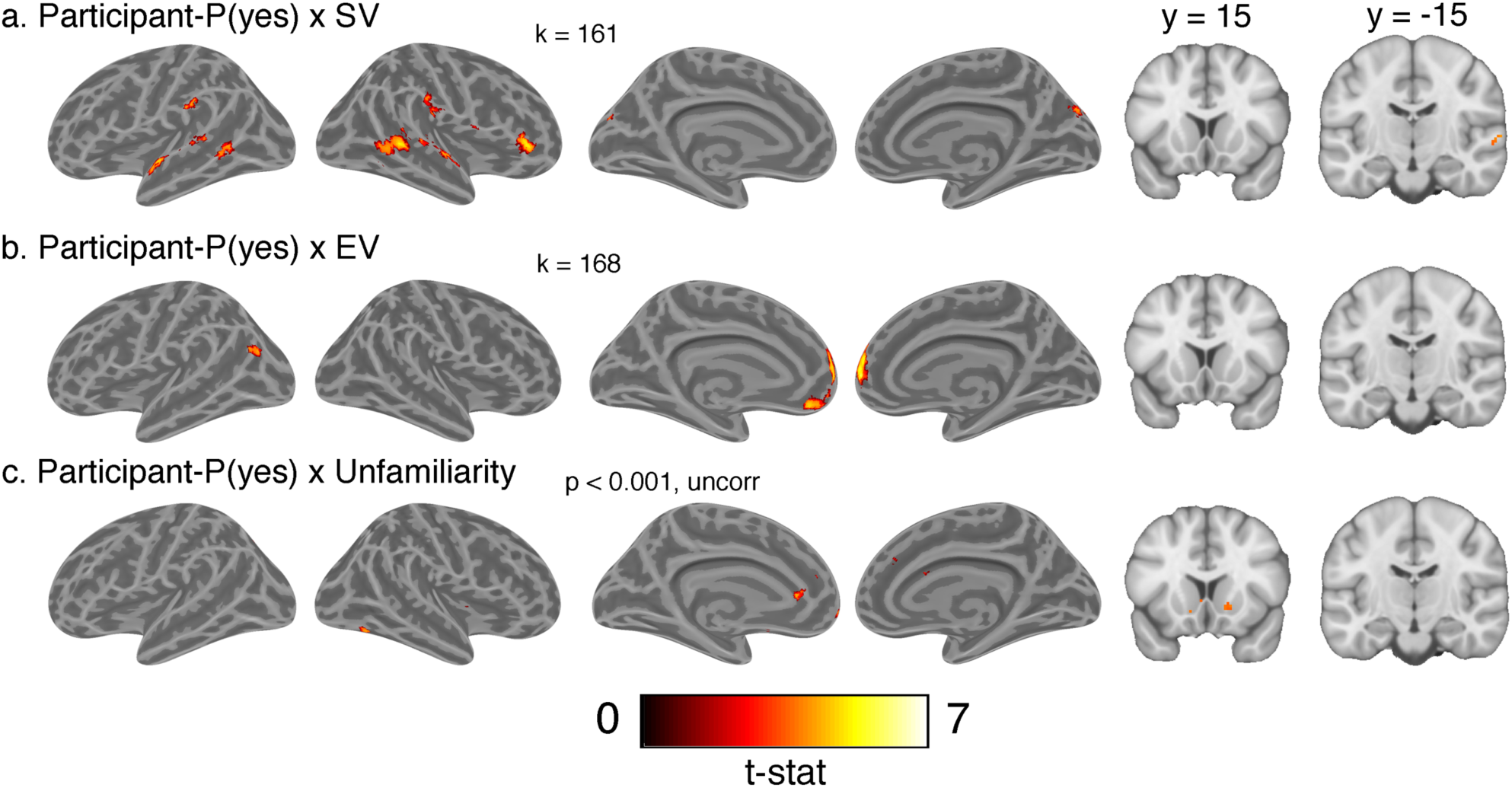
Whole brain t-statistic maps for interactions of model-based terms with participant-P(yes). **a.**. Interaction of participant-P(yes) and schema value (SV), **b.** Interaction of participant-P(yes) and experienced value (EV), **c**. Interaction of participant-P(yes) and unfamiliarity. All images are thresholded at *P* < 0.05, FWE cluster corrected, except for those shown in **c**. Cluster extent threshold for each map is given by value of k.

We next specifically compared how activity related to participant-P(yes) in vmPFC and striatum, the two regions we identified *a priori* as being associated with valuation, was modulated by experienced value, schema value and unfamiliarity. We extracted the mean beta coefficients for the interaction between participant-P(yes) and these three terms in the vmPFC and striatum ROIs. The interaction of participant-P(yes) with experienced and schema value was significant in vmPFC (one-tailed, one sample t-tests against zero: *t*(22) = 2.38, *P* ≤ 0.01, *d*’s ≥ 0.50). There was also a significant effect for the interaction of schema value with participant-P(yes) in the striatum (*t*(22) = 2.05, *P* = 0.03, *d* = 0.43), but not for the interaction of participant-P(yes) with experienced value. Further, the interaction of participant-P(yes) with unfamiliarity was significant in the striatum (*t*(22) = 2.30, *P* = 0.02, *d* = 0.48), but not vmPFC. Thus, vmPFC responded to greater agreement between both experienced and schema values with participant-P(yes), while striatum activity increased in response to participant-P(yes) when unfamiliarity was high, as well as the agreement of schema values with participant-P(yes).

To examine whether ROIs differed in their response to these value-related terms, we then tested an ROI (striatum, vmPFC) by modulator (interactions of participant-P(yes) with experienced value, schema value and unfamiliarity) interaction on these average beta coefficients (Figure 9). This interaction was highly significant (F _2,44_ = 8.67, *P* = 0.007, 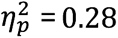), as was a main effect of ROI (F _1,22_ = 10.59, *P* = 0.004, 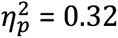), but not modulator (F _2,44_ = 0.10, *P* = 09, 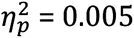). To determine the source of this interaction, we compared the difference of the betas in the vmPFC and striatum for each modulator (i.e. the effect of each interaction between ROIs). This difference was significantly greater for the interaction of participant-P(yes) with experienced value compared to both schema value and unfamiliarity (Bonferroni corrected t-tests *t*’s(22) ≥ 3.09, *P*’s < 0.05, *d* ≥ 0.64), but there was no comparable significant difference between schema value and unfamiliarity.

**Figure 9.**
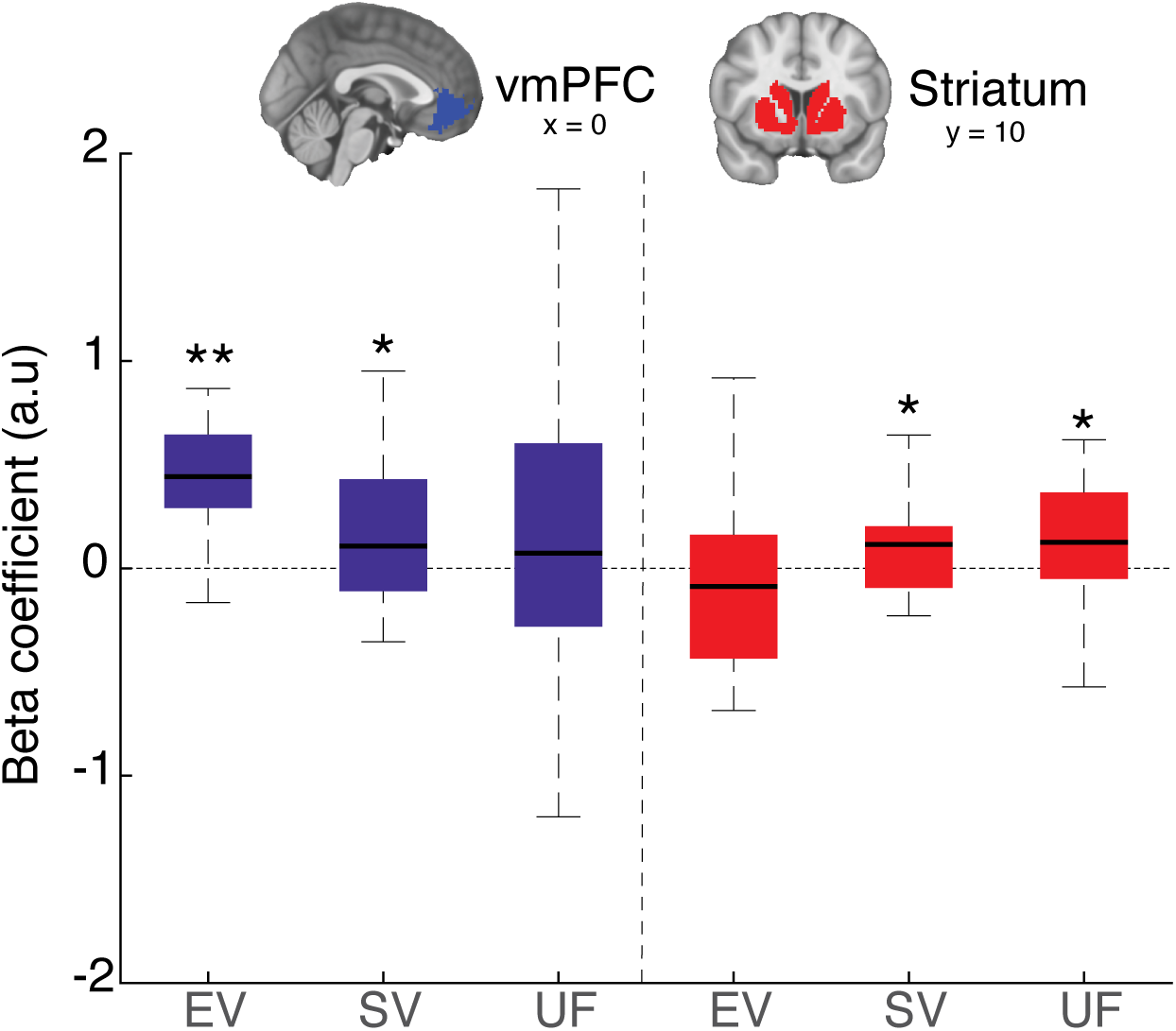
Boxplots showing mean beta coefficients from voxels in ventromedial prefrontal cortex (vmPFC) and striatum ROIs for interactions of participant-P(yes) with experienced value (EV), schema value (SV) and unfamiliarity (UF). ** *P* < 0.01, * *P* < 0.05, one-tailed one-sample t-test against zero.

To summarize, vmPFC activity reflected the influence of experienced and schema values on participants’ value judgments, and preferentially reflected the influence of explicitly experienced values on participants’ value judgments compared to the striatum. Striatal activity reflected the effect of unfamiliarity on participants’ value judgments and values inferred from a schema, but not value information stored directly in memory.

### Generalized psychophysical interactions

We hypothesized that differential demand on memory systems during decision-making would also lead to distinct patterns of functional connectivity, reflecting the contributions of different memory stores to the assessment of options. vmPFC has reciprocal connections with regions of middle temporal cortex (MTC) and anterior ventrolateral prefrontal cortex (aVLPFC) (Price, 2007; Saleem, Kondo, & Price, 2008). Both vmPFC and striatum also receive input from the hippocampus (HPC). We were principally interested in whether vmPFC and MTC connectivity increased with greater demands on schema-based decision-making, and whether striatum and HPC connectivity increased during the assessment of directly experienced values. To address this hypothesis, we carried out generalized psychophysical interaction (gPPI) analyses that contrasted how connectivity between pairs of ROIs listed in Table 1 changed between task conditions, as well as more exploratory seed-to-voxel whole-brain gPPI analyses focused on vmPFC, MTC, HPC and striatum.

We first examined whether connectivity with vmPFC and MTC was modulated by the schema-based value of ingredients on correct responses in the first presentation – i.e. when these values are correctly inferred from schematic knowledge and not retrieved from experience. Within an MTC seed, there was increased connectivity with vmPFC on the first presentation of positive recipe ingredients compared to negative recipe ingredients (Bonferroni corrected t-test: *t*(21) = 4.00, *P* < 0.01, *d* = 0.85), but not any other ROI (*P* ≥ 0.2, Bonferroni corrected). Similarly, using vmPFC as a seed ROI revealed increased connectivity with the MTC ROI in the same contrast (Bonferroni corrected t-test: *t*(21) = 3.68, *P* < 0.05, *d* = 0.78), but not any other ROI. An exploratory whole-brain seed-to-voxel analysis on this contrast with the MTC ROI as a seed found that this region had significantly greater connectivity with a cluster on the medial frontal wall overlapping with our vmPFC ROI (Figure 10a, MNI: −2, 46, −12), and a second cluster in lateral occipital cortex (MNI: −52, −68, 12). The same whole-brain ROI-to-voxel contrast with the vmPFC ROI as a seed did not reveal any significant clusters at an FWE cluster-corrected threshold.

**Figure 10.**
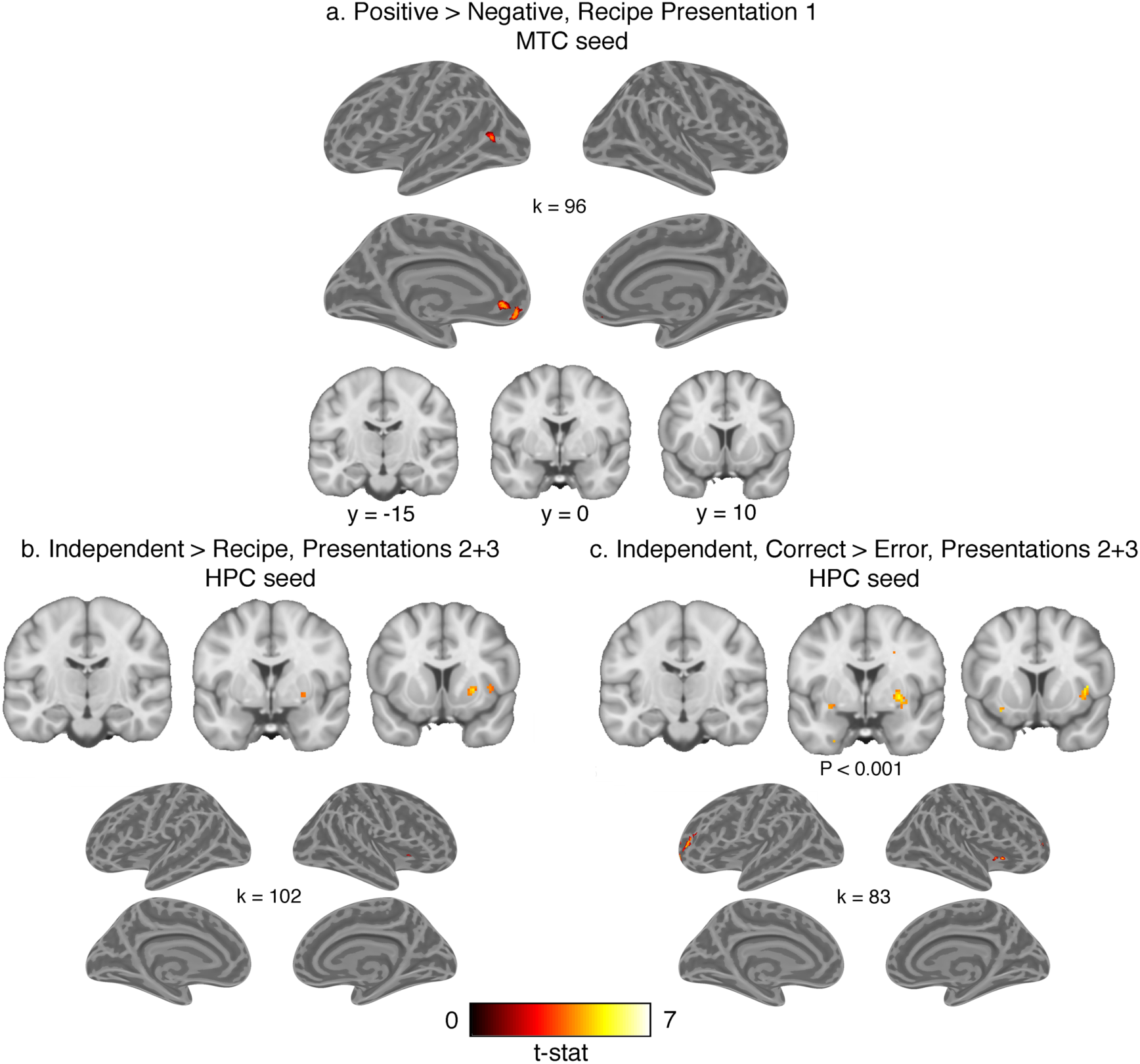
Whole brain t-statistic maps for seed-to-voxel generalized psychophysical interaction analyses. **a.** Voxels with increased connectivity with the middle temporal cortex (MTC) seed on the first presentation of positive recipe trials compared to negative recipe trials. **b.** Voxels with greater connectivity with the hippocampus (HPC) seed on the second and third presentation of independent trials compared to recipe trials. **c.** Voxels with increased connectivity with HPC on correct compared to error trials on the second and third presentations of independent ingredients. Voxels passing the *P* <0.05 FWE cluster corrected threshold are shown in all panels except for top row of **c.** Cluster extent threshold is given by k.

These data provide evidence that vmPFC and MTC connectivity changed depending on the value of recipe ingredients on the first presentation. To examine whether this effect might depend on successful retrieval of relevant schema-value information, we tested the interaction of value and accuracy on the first presentation of recipe ingredients. This interaction was significant when either vmPFC and MTC was used as a seed (Bonferroni corrected t-tests: *t*’s(20) ≥ 2.95, *P* < 0.05, *d* ≥ 0.64), but not between these seeds and any other ROIs. There was no main effect of accuracy on connectivity between ROIs on these trials. We further examined whether this value related increase in vmPFC-MTC connectivity was higher on the first presentation than subsequent presentations by testing the interaction of presentation number and value for recipe ingredients, but this interaction did not reach significance after correction for multiple comparisons in either seed.

Thus, vmPFC-MTC connectivity increased in response to retrieval of positive values inferred from schematic knowledge on this first presentation, and though this increase was greater on correct trials compared to errors, there were no main effects of clustering condition or accuracy. These null effects could partially stem from participants attempting to retrieve schematic associations on the first presentation in all conditions, consistent with greater MTC activity in these presentations compared to repetitions in the independent condition.

We also tested whether connectivity between MTC and vmPFC and any other ROIs depended on clustering status by contrasting the recipe and independent conditions on the first presentation, collapsing across value. There was no significant change in connectivity between MTC, vmPFC and any other ROIs in this contrast. Exploratory seed-to-voxel analyses for both of these contrasts were similarly null.

Next, we investigated if connectivity between with the HPC and striatum and other ROIs depended on the value of options in the second and third presentations. Contrasting positive and negative independent ingredients in these trials, we did not find any significant increase in connectivity between HPC and striatum with each other or any other region in ROI-to-ROI, or seed-to-voxel analyses. We further examined if connectivity with these ROIs was higher for positive than negative ingredients in the independent condition relative to the recipe condition by testing the interaction of value and clustering. This analysis was similarly null for both seeds.

We also tested if HPC and striatum connectivity depended on demands for retrieving previously experienced feedback associated with particular ingredient instance. Collapsing across value conditions, we contrasted independent and recipe ingredients on the second and third presentations where ingredient values could be retrieved from episodic memory, with the expectation that demand on episodic memory would be greater in the independent condition. There was no significant difference in connectivity between the striatum or HPC with any other ROIs in this contrast. However, a seed-to-voxel whole-brain analysis with the HPC seed showed significantly greater connectivity with a cluster in the right putamen and insula (Figure 10b; MNI: 26, 10, 0). The reverse seed-to-voxel analysis with the striatum ROI as a seed did not reveal significant clusters within a small-volume corrected HPC ROI.

To test if this increase in HPC and hippocampus connectivity was related to retrieval success we also contrasted correct and error responses on the second and third presentation of independent ingredients. Using the HPC as a seed in a seed-to-voxel analysis, we found increased connectivity with this ROI and the right putamen (MNI: 28, 2, 4) after small volume correction within the striatum ROI. An exploratory whole-brain analysis also found that HPC significantly increased connectivity with clusters in bilateral anterior rostrolateral PFC and right insula (Figure 10c). However, there were no significant differences in connectivity between these conditions in ROI-to-ROI contrasts with either HPC or striatum as seeds. The reverse ROI-to-voxel analysis with the striatum ROI as a seed did not reveal significant increase in connectivity in a small-volume corrected HPC ROI. All peaks from seed-to-voxel analyses are reported in Table 5.

**Table 5.** Whole brain seed-to-voxel connectivity effects from Figure 10

| Region (AAL2) | MNI Coordinates |  |  | Number of voxels | Peak *t*-value |
| --- | --- | --- | --- | --- | --- |
|  | x | y | z |  |  |
| **Recipe Positive > Recipe negative, presentation 1, correct — MTC seed (k = 96)** |  |  |  |  |  |
| Left superior frontal gyrus, medial orbital | -2 | 46 | -12 | 351 | 6.42 |
| Left middle temporal gyrus | -52 | -68 | 12 | 112 | 5.30 |
| **Correct > Error, independent presentations 2 and 3, correct — HPC seed (k = 83)** |  |  |  |  |  |
| Left middle frontal gyrus | -40 | 52 | 20 | 253 | 6.14 |
| Left superior frontal gyrus, dorsolateral | -18 | 54 | -4 | 253 | 5.66 |
| Right insula | 36 | 18 | 0 | 100 | 5.76 |
| Right superior frontal gyrus, dorsolateral | 30 | 54 | 14 | 122 | 5.76 |

All reported clusters were significant at the $P < 0.05$ , FWE-corrected threshold at the cluster level after peak thresholding at $P < 0.001$ . The critical cluster extent threshold for each contrast is given by the value of *k*.

In summary, we found that vmPFC-MTC connectivity depended on recipe value associations on the first presentation where this information was key to guiding choice. Meanwhile, HPC increased connectivity with the right putamen in response to greater demand on episodic memory compared to trials where schema memory could still be used, and trials where value information was not successfully retrieved.

## Discussion

Here we investigated the neural circuitry involved in value judgments based on distinct memory sources. We had put forward three main hypotheses: (1) vmPFC is preferentially involved in inferred value judgments based on schematic world knowledge, (2) striatum is preferentially involved in representing experienced values, (3) vmPFC and striatum interact with distinct memory networks in line with their respective involvement in building values from schematic knowledge and experience.

Our results were inconsistent with these *a priori* hypotheses, but yielded new insights into the brain systems linking memory with value-based decision-making. Striatum responded to the effect of the unfamiliarity of options on participants’ value judgments and schema-based values. In contrast, vmPFC responded to the influence of value judgments based both on direct experience and schema memory. Analysis of connectivity between these regions and memory stores revealed greater connectivity between the striatum and HPC in response to demands on recall of experienced values, while vmPFC and MTC connectivity increased in response to values derived from schema memory. Thus, vmPFC represented retrieved value information from multiple memory sources, and coordinated with MTC when value was inferred. In contrast, the striatum incorporated basic memory strength into the decision process, and increased connectivity with HPC in response to demand for retrieval of experienced values.

Participants were able to infer the value of ingredients based on schematic knowledge without feedback, and continued showing a benefit from recipe information in subsequent presentations. Our computational model captured this behavior using normative information about ingredient-recipe associations. Notably, freely fitting the number of dimensions used in reconstructing these recipe-ingredient associations improved the fit of the model, implying that participants varied in the depth of semantic knowledge they employed in this task. The positive recipe also had a greater influence on behavior compared to the negative recipe, indicating that participants prioritized searching associations with this recipe first — possibly because they adopted a default response to pass on ingredients where they could not retrieve positive value information. Variance in how participants engage with these schemas to form value estimates may also be reflected in neural activity, however we did not have a sufficient sample size to make a strong test of such individual differences.

Memory strength, expressed as unfamiliarity, had a substantial impact on participants’ willingness to endorse ingredients. Participants demonstrated a general tendency against betting on independent ingredients on the first presentation and against betting on ingredients where memory strength was lower on subsequent presentations. Given that there was an equal proportion of positive and negative value ingredients, and a 2:1 ratio of points gained to lost on a bet, this behavior reflects a deviation from a rational maximization of expected value, akin to loss aversion (Kahneman & Tversky, 1979). While the proportion of positive to negative value trials was not made explicit to the participants, and we cannot be sure that participants were aware of this equal weighting, this proportion was the same in the practice phase and main experiment, so that participants would have had experience with an equal proportion of positive and negative trials before starting the main task. Whereas uncertainty is often operationalized as descriptive or experienced probabilities of outcomes in behavioral economics, here it depended on the strength of participants’ memory of outcomes. These results are also consistent with work showing a bias toward remembered items in preference-based choice (Gluth, Sommer, Rieskamp, & Buchel, 2015). In real terms, people appear to apply a meta-cognitive strategy where the strength of evidence from memory affects option value, regardless of its content. We would thus predict that a strong memory for a weakly positive or negative value would be more likely to drive a response than a weak memory for a strongly positive or negative value.

Our results indicate that vmPFC represents values based on multiple memory sources. These findings add to past work demonstrating that the value function represented in vmPFC can flexibly bend to accommodate attentional priority toward different option attributes (Hare, Malmaud, & Rangel, 2011; Hunt, Dolan, & Behrens, 2014; Leong, Radulescu, Daniel, DeWoskin, & Niv, 2017; Rudorf & Hare, 2014). However, we cannot be sure if the values represented here are ultimately referenced to the points earned by the participant, or reflect a social value based on the experience of the “customer” whom they are ostensibly feeding according to the cover story for the task.

We had predicted that vmPFC would be engaged more by schema-based value inference that retrieval of a specific experience. This prediction was based on lesion evidence demonstrating that vmPFC is not necessary for value judgment *per se* (Reber et al., 2017; Vaidya & Fellows, 2015; Yu, Dana, & Kable, 2018), but is needed for incorporating higher-order information into value judgments (Vaidya, Sefranek, & Fellows, 2017; Xia, Stolle, Gidengil, & Fellows, 2015). Other work has shown that inactivation or damage to vmPFC or OFC does not affect decision-making or value judgment in situations where values can be retrieved from prior direct experience, but does so when this information must be inferred from structured knowledge about a task (Bradfield et al., 2015; Izquierdo et al., 2004; Jones et al., 2012; Rudebeck, Saunders, Prescott, Chau, & Murray, 2013).

There may be a few factors contributing to this discrepancy. While experienced values were learned over many repeated exposures in these lesion experiments, here they were learned from just one or two trials. Recent work has shown that spaced, brief training over a period of days in a reinforcement learning task increases fMRI value signals in vmPFC compared to an equivalent number of trials within a single session (Wimmer, Li, Gorgolewski, & Poldrack, 2018), implying that this region is engaged more when value judgments rely on episodic memory rather than massed experience. Although our results suggest the involvement of vmPFC in representing both directly experienced and schema-based values, this information may be robustly represented by other regions that do not depend on vmPFC to influence behavior. Studying the effects of vmPFC damage in this task could help assess when this region is critical for memory-based value judgment.

Our hypotheses are not fully committed to the precise format of experienced values in this task. Our design required that participants store information about feedback paired with ingredients in a small number of trials as a specific experience separated by other trials over a long interval. We assumed that participants would mostly rely on episodic memory for robust retention of such information. However, we cannot rule out the possibility that value information would also be stored in other ways (e.g. in working memory, or through conditioned stimulus-response mappings). Nonetheless, this possibility does not change our ability to contrast inference of values from a memory schema and reliance on direct experience more generally.

Notably, our results indicate that interactions between vmPFC and MTC support inference of values from semantic knowledge. Other work has shown that utilization of semantic meaning in value judgments increases interactions between vmPFC and regions of left middle and superior temporal cortex (Lim, O’Doherty, & Rangel, 2013). Here, vmPFC-MTC connectivity depended on the value of schematic associations. This result could reflect the accumulation of value information through the interaction between these regions, or the differential weight given to positive and negative recipes, as there was behavioral evidence that participants prioritized positive recipe associations. In either case, these data provide evidence that engagement between vmPFC and semantic memory stores is involved in inferential value judgments based on real-world structured knowledge in absence of explicit experienced values.

This experiment explicitly crossed two features commonly associated with the vmPFC, but rarely compared within the same task: value coding and schema-based memory and inference (Chan, Niv, & Norman, 2016; Delgado et al., 2016; Ghosh, Moscovitch, Melo Colella, & Gilboa, 2014). Our data are consistent with both proposed roles for this region: with schema-related knowledge playing a role in value construction in vmPFC, and with this region responding to utilization of schema-related knowledge more generally. Recent rodent electrophysiological data also indicate that value and schematic task representations co-exist in separable codes within OFC (Zhou et al., 2019), suggesting these functions may co-exist within the same regions.

The role of striatum in memory tasks has been the focus of recent research, with several hypotheses being considered (Elward et al., 2015; King, Chastelaine, Elward, Wang, & Rugg, 2018; Schwarze, Bingel, Badre, & Sommer, 2013; Scimeca & Badre, 2012). Here, activity in the striatum integrated the influence of memory strength on participants’ decisions to bet or pass on ingredients, as its activity correlated with the interaction of participant-P(yes) and unfamiliarity. Relatedly, recent work from our group has observed that the striatum tracked prediction errors for retrieval success during a recognition memory task (Scimeca et al., 2016), analogous to its role in action selection and reinforcement-learning (Garrison, Erdeniz, & Done, 2013; Haber & Knutson, 2010; Pessiglione, Seymour, Flandin, Dolan, & Frith, 2006). These prediction errors were correlated with changes in the criterion by which evidence from memory was evaluated. Unlike conventional reinforcement learning settings, the striatum was sensitive to prediction errors that derived from memory strength rather than the value of a response. We expand on those findings here, demonstrating that the striatum responds to the influence of memory strength on participants’ willingness to make a bet in a value-based decision-making task, independent of memory valence. Together, these results imply a role for the striatum in a cognitive decision about when to act based on memory strength, equivalent to its role in controlling a tendency to bet in risky decision-making (Knutson, Wimmer, Kuhnen, & Winkielman, 2008; Nachev et al., 2015; Zalocusky et al., 2016), or in gating working memory (Chatham & Badre, 2015).

Notably, the striatum responded to the strength of memory for experienced values rather than the experienced values of options themselves. The striatum responds to successful memory retrieval in many tasks (Spaniol et al., 2009), though the functional significance of this activation is debated. Some work has argued that these signals reflect a value signal that depends on the congruence of the memory task with participants’ goals (Han et al., 2010), while others have suggested that striatal reward and retrieval signals are distinct (Elward et al., 2015). Here we found evidence that unsigned memory strength (i.e. unfamiliarity) modulates striatal value signals, but the signed experienced value of options did not. These data indicate that value and retrieval signals in the striatum may follow a common dimension that corresponds to the utility of retrieved information given task demands — biasing cortical representations of associated actions based on the strength of information recovered from retrieval.

We also found that the striatum responded to the interaction of participant-P(yes) and schema value. While of a smaller effect size, these data imply that the striatum encodes these inferred values in addition to the effects of unfamiliarity. Indeed, previous work has found that the striatum responds to prediction errors inferred structured knowledge about a task (Daw, Gershman, Seymour, Dayan, & Dolan, 2011; Lohrenz, McCabe, Camerer, & Montague, 2007).

We also observed that connectivity between HPC and the striatum increased in the second and third presentations of independent ingredients compared to recipe ingredients, and on correct choices for independent ingredients compared to error trials. This result implies that information shared in a HPC-striatal circuit is involved in successful retrieval of experienced ingredient-value associations. While this result depended on the choice of seed ROI, this apparent selectivity might owe to our striatal ROI averaging over functionally distinct subregions.

In conclusion, we demonstrated the involvement of the striatum and vmPFC in distinct processes within memory-based decision-making. Our data provide fresh insights into the dynamics of this circuitry, with a view into the intersection of decision-making and memory that is the seat of much real-world choice behavior. These results have implications for understanding the origins of decision-making impairments in neurodegenerative diseases such as Alzheimer’s disease and frontotemporal dementia (Salmon & Bondi, 2009), as well as Parkinson’s disease (Edelstyn, Shepherd, Mayes, Sherman, & Ellis, 2010). New insights into the interaction of these circuits may have relevance for clinical interventions or treatments.

## Conflicts of interest

The authors declare no competing conflicts of interest.

## Acknowledgements

This work was supported by a Multidisciplinary University Research Initiative award from the Office of Naval Research (N00014-16-1-2832), an award from the James S. McDonnell Foundation, and a Ruth L. Kirschstein National Research Service Award to A.V. (NIMH F32 MH116592-01A1). We are grateful to Michael Frank for helpful comments and suggestions on the manuscript, as well as Apoorva Bhandari and Andra Geana for constructive discussion on methods and analysis. Thanks also to Adriane Spiro for help with data collection, and Henry Jones for assistance with behavioral piloting and analysis of normative data.

## Notes

### Competing Interest Statement

The authors have declared no competing interest.

### Summary of Updates

Computational modeling approach has been revised (cross-validation is now used for model comparison rather than AIC), as have model-based fMRI analyses. Several smaller changes have been made throughout the manuscript for better clarity.

